# Stochastic Misfolding Drives the Emergence of Distinct α-Synuclein Strains

**DOI:** 10.1101/2025.06.03.657690

**Authors:** Raphaella W.L. So, Benedikt Frieg, José D. Camino, Nicholas R.G. Silver, Alison Mao, Erica Stuart, Gunnar F. Schröder, Joel C. Watts

## Abstract

The existence of α-synuclein conformational strains provides a potential explanation for the clinical and pathological differences among synucleinopathies such as Parkinson’s disease and multiple system atrophy. However, how distinct α-synuclein strains are formed *in vivo* remains unknown. Here, we examined whether unique strains of self-propagating α-synuclein aggregates can arise within a consistent molecular environment. Unexpectedly, we observed conformational heterogeneity between individual preparations of α-synuclein pre-formed fibrils (PFFs) generated by polymerizing recombinant wild-type or A53T-mutant human α-synuclein under identical conditions. Moreover, we found that α-synuclein aggregates formed spontaneously in the brains of a transgenic synucleinopathy mouse model were conformationally diverse, leading to the identification of three distinct disease subtypes. Propagation of putative PFF- and brain-derived α-synuclein strains in mice initiated several distinct synucleinopathies, characterized by differences in disease onset times, cerebral α-synuclein deposition patterns, and the conformational attributes of α-synuclein aggregates. The conformational diversity of α-synuclein aggregates across PFF preparations and between the brains of individual transgenic mice demonstrates that α-synuclein can spontaneously form multiple self-propagating strains within an identical environment both *in vitro* and *in vivo*. This suggests that stochastic misfolding into distinct aggregate structures drives the emergence of α-synuclein strains and implies that the intrinsic variability of common synucleinopathy research tools must be considered when designing and interpreting experiments.

## Introduction

α-Synuclein (α-syn) is a 140-amino-acid intrinsically disordered cytoplasmic protein found at the presynaptic nerve terminal that can adopt an α-helical conformation upon binding to membranes [1–3]. Under physiological conditions, α-syn participates in SNARE complex assembly and mediates vesicular membrane fusion [4]. During disease, α-syn assembles into insoluble, β-sheet-rich protein deposits that are highly phosphorylated at serine-129 (PSyn) [5, 6]. The presence of α-syn aggregates in the central nervous system is a pathological hallmark of the “synucleinopathies”, which include progressive neurodegenerative diseases such as Parkinson’s disease (PD), dementia with Lewy bodies (DLB), and multiple system atrophy (MSA) [7, 8]. The brains of PD and DLB patients are characterized by neuronal α-syn inclusions in the form of Lewy bodies (LB) and Lewy neurites, whereas the brains of MSA patients predominantly exhibit glial cytoplasmic α-syn inclusions in oligodendrocytes. α-Syn plays a critical role in synucleinopathy pathogenesis since polymorphisms within the α-syn-coding gene, *SNCA*, have been identified as risk factors for sporadic PD [9, 10], and multiplication or mutation of *SNCA* causes familial forms of PD and DLB [11–17]. One of the most widely studied α-syn mutations is A53T, which causes early-onset PD [13]. In transgenic mice, overexpression of A53T-mutant human α-syn results in progressive motor dysfunction accompanied by a cerebral synucleinopathy [18].

Disease progression in the synucleinopathies is believed to be caused by the cell-to-cell propagation of α-syn aggregates within the brain [19, 20]. Propagation of α-syn aggregates is thought to occur by a “prion-like” mechanism in which pre-existing aggregates act as a template or “seed” to the direct the misfolding of normal α-syn into a growing fibrillar structure [21]. As α-syn is a cytoplasmic protein, cellular spreading likely involves several steps, including fragmentation of α-syn aggregates into smaller seeds, transfer of an α-syn seed from one cell to another, and seeding of normal α-syn in the recipient cell to initiate new aggregate formation. α-Syn propagation can be studied experimentally by injecting non-transgenic or transgenic mice expressing wild-type (WT) or mutant α-syn with pre-formed α-syn aggregates and monitoring the deposition and spread of cerebral α-syn pathology over time [22–29]. Recombinant α-syn can be polymerized into aggregates by shaking at 37 °C *in vitro*, and these pre-formed fibrils (PFFs) are effective at inducing the pathological accumulation of α-syn in cultured cells and mice [23, 30, 31]. Brain extracts from PD, DLB, and MSA patients can also be used to seed α-syn aggregation in cultured cells and mice, although α-syn aggregates from MSA brains are more potent at inducing α-syn deposition [25, 26, 32–35].

As *SNCA* mutations account for only a small proportion of total synucleinopathy cases, inclusions composed of WT human α-syn are present in most PD and DLB patients as well as in all MSA cases. One potential explanation for how a single protein can cause several distinct diseases is the conformational strains hypothesis. This theory, which originated in the prion diseases, posits that α-syn can misfold in several different ways to generate structurally distinct aggregates, each of which causes a different synucleinopathy [36, 37]. Indeed, cryo-electron microscopy (cryo-EM) studies have revealed that α-syn aggregates isolated from the brains of patients with different synucleinopathies are structurally dissimilar [38–40]. Upon injection of α-syn aggregates into laboratory animals, conformational strains can be distinguished by differences in clinical symptoms, rates of disease progression, neuropathological profiles, and the biochemical properties of the resultant α-syn aggregates [36]. Due to the process of templated conformational conversion, true α-syn strains should also be able to undergo repeated (serial) passaging in an animal model without a change in their conformational properties.

Considerable evidence for the conformational strains hypothesis has come from *in vitro* studies using recombinant α-syn aggregates. Structurally distinct preparations of α-syn PFFs are often referred to as “polymorphs” rather than strains since their propagation properties may not yet have been assessed *in vivo*. Distinct polymorphs of α-syn PFFs can be generated by changing the chemical conditions during aggregation. For example, human WT α-syn PFFs formed in the presence or absence of KCl differ in their relative toxicities and ability to seed PSyn pathology in rodent models [41–43]. Moreover, serial propagation of α-syn PFFs *in vitro* led to the emergence of two distinct α-syn polymorphs with a differential ability to induce either α-syn or tau aggregation in neurons [44–46]. Finally, polymerization of recombinant human α-syn containing the A53T mutation in the presence or absence of NaCl generated two distinct α-syn polymorphs that propagated as different strains upon inoculation into transgenic mice that express sequence-matched α-syn [47]. Thus, α-syn can assemble into multiple distinct strains of self-propagating aggregates.

While buffer composition can influence α-syn aggregation *in vitro*, how distinct α-syn strains arise naturally *in vivo* remains unknown. One study found that distinct cellular milieus present within different brain cell types influence the propagation properties of α-syn aggregates, with passage of misfolded α-syn through oligodendrocytes, but not neurons, leading to production of a more aggressive strain [48]. Another study found that α-syn polymorphs with differential reactivities to an amyloid-binding dye can emerge spontaneously upon polymerization of recombinant WT human α-syn *in vitro* [49]. Here, we investigated whether unique strains of self-propagating α-syn aggregates can form under identical molecular conditions. Using both PFFs composed of either WT or A53T-mutant human α-syn as well as transgenic mice that develop a spontaneous synucleinopathy, we found that the same molecular environment can give rise to multiple α-syn strains that initiate biochemically and neuropathologically distinct synucleinopathies upon propagation in a mouse model. We therefore conclude that an initial stochastic misfolding event drives the formation of different α-syn conformational strains that then propagate via templated conformational conversion.

## Results

### Recombinant human A53T-mutant α-syn assembles into two distinct polymorphs

Previously, we showed that polymerization of recombinant A53T-mutant human α-syn in either the presence or absence of 100 mM NaCl led to the respective generation of “Salt (S)” and “No Salt (NS)” PFFs that gave rise to unique disease phenotypes upon inoculation into hemizygous M83 transgenic mice (M83^+/-^) [47]. In follow-up experiments, analysis of a larger number of S PFF preparations led to the unexpected discovery that different α-syn fibril polymorphs can be discerned amongst experimental replicates. To address this issue systematically, we generated 30 independent preparations of A53T-mutant α-syn PFFs formed under identical conditions by continuous shaking for 7-14 days in a buffer containing 100 mM NaCl (**Fig. 1a**). The conformational properties of the individual A53T PFF preparations were characterized by digestion with proteinase K (PK), a technique we previously used to discriminate between S and NS PFFs [47]. Based on the pattern of PK-resistant α-syn fragments observed following digestion, two distinct polymorphs of A53T PFFs could be identified, and these were termed A53T_1_ PFFs and A53T_2_ PFFs. A53T_1_ PFFs were characterized by two dominant PK-resistant bands at ∼11 and ∼9 kDa whereas A53T_2_ PFFs were characterized by a single dominant band at ∼9 kDa (**Fig. 1b**). The two PFF types occurred at similar frequencies (**Fig. 1c**) and were both observed when PFFs were generated from two independent batches of recombinant A53T-mutant human α-syn (**Supplemental Fig. 1a**).

**Figure 1.**
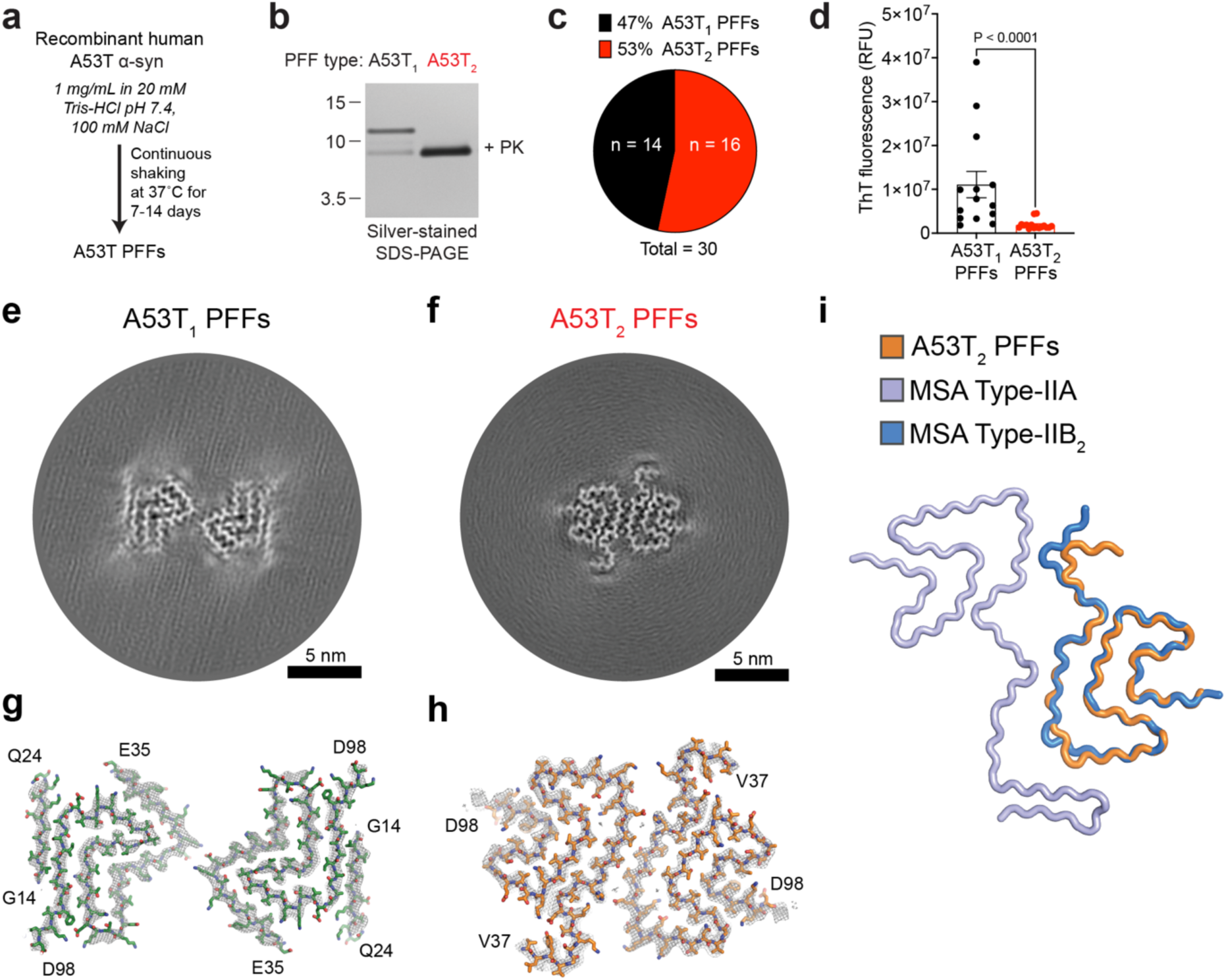
A53T-mutant recombinant α-syn forms two distinct PFF polymorphs. **a**) Schematic for formation of A53T PFFs. **b**) Following PK digestion of A53T PFFs and analysis by SDS-PAGE, differences in the pattern of PK-resistant insoluble α-syn fragments reveals two distinct PFF polymorphs (A53T_1_ and A53T_2_). Molecular weight markers indicate kDa. **c**) Relative frequencies of A53T_1_ and A53T_2_ polymorphs across 30 independent PFF preparations. **d**) ThT fluorescence (mean ± s.e.m.) of PFF preparations classified as either A53T_1_ (n = 14) or A53T_2_ (n = 16). Statistical significance was assessed using a Mann-Whitney test. **e, f**) Cross-sections of the cryo-EM density maps showing A53T_1_ (e) and A53T_2_ (f) PFFs. Scale bars indicate 5 nm. **g, h**) Overlay of sharpened high-resolution cryo-EM density maps (gray mesh representation) and the atomic models of the A53T_1_ (g) and A53T_2_ (h) PFFs. **i**) Superimposition of the A53T_2_ protofilament structure (orange) with an α-syn Type II-2 filament extracted from brain tissue of individuals with MSA (PDB: 6XYQ). The IIA protofilament is colored in light purple and the IIB_2_ protofilament is colored in blue.

After assigning each PFF preparation as either A53T_1_ or A53T_2_ using PK digestion, we performed additional characterization of the two polymorphs. A53T_1_ PFFs could be distinguished from A53T_2_ PFFs by significantly higher fluorescence following reaction with the protein aggregate-binding dye Thioflavin T (ThT) (**Fig. 1d**). While this could imply differences in the quantity of fibrils formed, distinct ThT binding efficiencies can also indicate conformational differences between α-syn aggregates [50]. Previously, we used a curcumin dye-binding assay as well as a conformational stability assay that measures the resistance of α-syn aggregates to denaturation by guanidine hydrochloride (GdnHCl) to differentiate between S and NS PFFs [47]. However, we found that neither assay was capable of distinguishing between the A53T_1_ and A53T_2_ polymorphs (**Supplemental Fig. 1b-f**). Abundant fibrils were observed by negative stain electron microscopy in both A53T_1_ and A53T_2_ PFF preparations (**Supplemental Fig. 1g**). To conclusively determine whether the A53T_1_ and A53T_2_ PFFs are structurally distinct, we solved their 3D structures using cryo-EM (**Fig. 1e-h**). While both A53T_1_ and A53T_2_ PFFs were composed of two protofilaments, their individual protofilament structures differed considerably. Interestingly, the unstructured region between amino acids Q24 and E35, which disrupts the fold of A53T_1_ PFFs, may explain the presence of two major protease-resistant fragments following PK digestion (**Fig. 1b**). The structure of the A53T_2_ PFFs was very similar to that of Type-IIB_2_ α-syn protofilaments isolated from MSA patients (**Fig. 1i**) [39]. Collectively, these results demonstrate that two structurally distinct PFF polymorphs can emerge when A53T-mutant human α-syn is polymerized in a physiological buffer.

### A53T_1_ and A53T_2_ PFFs generate distinct α-syn strains upon propagation in M83^+/-^ mice

To assess the propagation behavior of the A53T_1_ and A53T_2_ polymorphs *in vivo*, we intracerebrally inoculated groups of M83^+/-^ mice with three independent preparations each of A53T_1_ or A53T_2_ PFFs (**Fig. 2a**). The preparations of A53T_1_ and A53T_2_ PFFs analyzed by cryo-EM were included in the replicates. As a negative control, mice were inoculated with monomeric (unpolymerized) recombinant A53T-mutant human α-syn. M83^+/-^ mice were chosen for the A53T PFF propagation studies as they 1) express sequence-matched A53T-mutant human α-syn [18]; 2) only rarely develop a spontaneous synucleinopathy at advanced ages; and 3) exhibit progressive neurological illness involving hind-limb paralysis, slowness of movement, loss of grip strength, and weight loss upon injection with α-syn PFFs or MSA brain extract [26, 47]. Digestion of the A53T PFF samples used as inocula with PK revealed the absence of PK-resistant α-syn species in the monomer control, while the three independent replicates of A53T_1_ and A53T_2_ PFFs demonstrated their expected patterns of PK-resistant α-syn (**Fig. 2b**). Neurological disease developed significantly more rapidly in M83^+/-^ mice inoculated with A53T_2_ PFFs than in mice inoculated with A53T_1_ PFFs (**Fig. 2c**). All 25 mice inoculated with A53T_2_ PFFs developed neurological illness whereas only 18 of the 23 mice inoculated with A53T_1_ PFFs became symptomatic (**Supplemental Table 1**). For the A53T_1_ replicates, the PFF preparation that exhibited the highest ThT fluorescence resulted in the most rapid and efficient transmission. Whereas 8 of 9 control mice inoculated with monomeric A53T α-syn remained asymptomatic at 540 days post-inoculation (DPI), one mouse developed neurological illness at 529 DPI (**Fig. 2c**), and the brain of this mouse displayed signs of synucleinopathy (**Supplemental Fig. 2**). This is consistent with the rare occurrence of late-onset spontaneous illness in aged M83^+/-^ mice [18].

**Figure 2.**
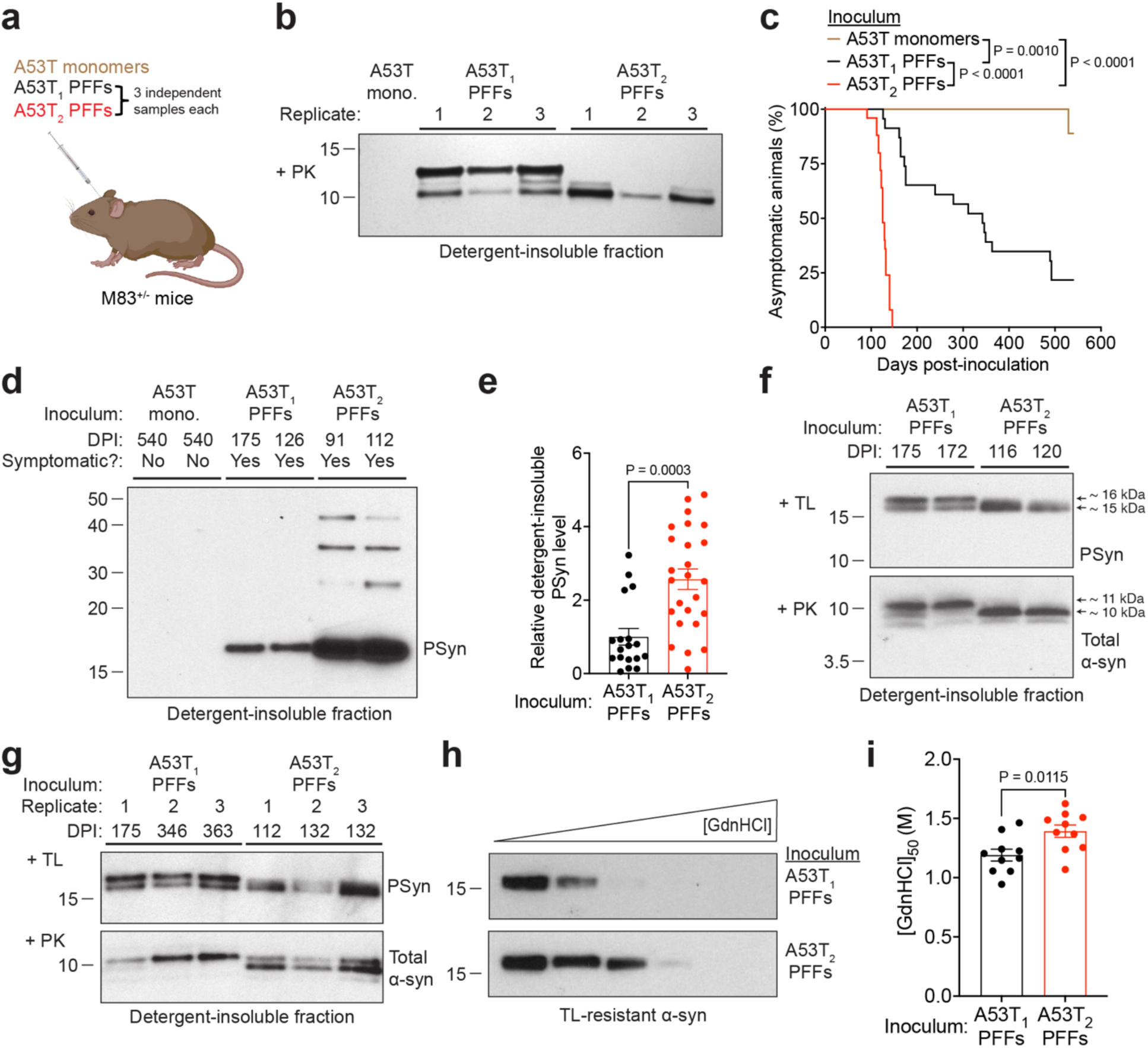
Propagation of A53T PFF polymorphs in M83^+/-^ mice. **a**) Schematic of intracerebral inoculation experiments in M83^+/-^ mice. **b**) Silver-stained SDS-PAGE of the detergent-insoluble fraction following PK digestion of monomeric (mono) recombinant A53T-mutant α-syn and the PFF samples used for inoculation experiments. **c**) Kaplan-Meier survival curves for M83^+/-^ mice inoculated with monomeric A53T-mutant α-syn (n = 9), A53T_1_ PFFs (n = 23), or A53T_2_ PFFs (n = 25). For the PFF inoculations, data from the three replicate experiments for each polymorph is pooled. Statistical significance was determined using the Log-rank test. **d**) Immunoblot of detergent-insoluble PSyn species in brain extracts from M83^+/-^ mice at the indicated DPI with either monomeric A53T α-syn (asymptomatic mice) or A53T PFFs (symptomatic mice). Two representative mice for each inoculum are shown. **e**) Quantification of detergent-insoluble PSyn species in brain extracts from symptomatic M83^+/-^ mice inoculated with either A53T_1_ PFFs (n = 18) or A53T_2_ PFFs (n = 25). **f**) Immunoblots of detergent-insoluble TL-resistant PSyn species and PK-resistant total α-syn species in brain extracts from symptomatic M83^+/-^ mice at the indicated DPI with A53T_1_ PFFs or A53T_2_ PFFs. Two representative mice for each inoculum are shown. **g**) Immunoblots of detergent-insoluble TL-resistant PSyn species and PK-resistant total α-syn species in brain extracts from symptomatic M83^+/-^ mice at the indicated DPI with individual replicates of A53T_1_ PFFs or A53T_2_ PFFs. **h**) Representative immunoblots for residual insoluble TL-resistant α-syn species following treatment of brain extracts from symptomatic M83^+/-^ mice inoculated with A53T_1_ or A53T_2_ PFFs with increasing concentrations of GdnHCl. **i**) [GdnHCl]_50_ values for TL-resistant α-syn species in symptomatic M83^+/-^ mice inoculated with either A53T_1_ or A53T_2_ PFFs (n = 9 each, 3 mice from each replicate experiment). In panels e and i, graphs display mean ± s.e.m. and statistical significance was assessed using a Mann-Whitney test. In panels b, d, f, g, and h, molecular weight markers indicate kDa.

Whereas detergent-insoluble α-syn species were observed in brain extracts from all M83^+/-^ mice regardless of inoculum, likely due to the overexpression of α-syn in the M83 line (**Supplemental Fig. 3a**), detergent-insoluble PSyn species indicative of pathological α-syn aggregation were present in symptomatic PFF-inoculated mice but absent in asymptomatic monomer-injected mice (**Fig. 2d**). Likewise, protease-resistant insoluble α-syn species were observed in brain extracts from symptomatic M83^+/-^ mice injected with A53T_1_ or A53T_2_ PFFs but not in extracts from mice that remained asymptomatic following injection with monomeric α-syn (**Supplemental Fig. 3b**). Detergent-insoluble PSyn levels were significantly higher in brain extracts from A53T_2_-injected mice than from A53T_1_-inoculated animals (**Fig. 2e**). In addition, limited proteolysis by digestion with thermolysin (TL) or PK revealed conformational differences between α-syn aggregates found in brain extracts from mice inoculated with the two PFF polymorphs. α-Syn aggregates induced by A53T_1_ PFFs were characterized by two equally intense TL-resistant bands at ∼16 and ∼15 kDa, as well as a dominant PK-resistant band at ∼11 kDa. In contrast, α-syn aggregates induced by A53T_2_ PFFs possessed a single ∼15 kDa TL-resistant band, as well as a dominant PK-resistant band at ∼10 kDa (**Fig. 2f**). The same protease digestion patterns were observed across mice inoculated with each of the three replicates of the same PFF polymorph (**Fig. 2g**). Using a conformational stability assay, we found that α-syn aggregates in brain extracts from A53T_2_ PFF-inoculated M83^+/-^ mice were more resistant to GdnHCl denaturation, which could indicate a more tightly packed structure (**Fig. 2h, i**).

Histological staining revealed significantly higher amounts of cerebral PSyn deposition in PFF-inoculated M83^+/-^ mice compared to mice inoculated with monomeric α-syn (**Supplemental Fig. 4**). Levels of PSyn deposition in the pons and midbrain were significantly higher in A53T_2_-inoculated mice than in A53T_1_-inoculated animals, while no difference was found in the hypothalamus (**Fig. 3a**). In addition, mice inoculated with A53T_2_ PFFs had more PSyn deposits in the cortex (**Fig. 3b, c**) and the dentate gyrus of the hippocampus (**Fig. 3d, e**) than mice inoculated with A53T_1_ PFFs. Previously, we reported that neurons within the midbrains of M83^+/-^ mice inoculated with either S or NS PFFs developed PSyn inclusions with distinct morphologies. “Ring-like” PSyn deposits were found in mice inoculated with S PFFs whereas “LB-like” PSyn inclusions were found in mice inoculated with NS PFFs [47, 51]. The PSyn aggregates in the midbrains of both A53T_1_- and A53T_2_-inoculated mice were predominantly ring-like (**Fig. 3f**), with no significant differences in morphology observed between the two groups of mice (**Fig. 3g**). Based on the unique disease incubation periods, distinct biochemical fingerprints of α-syn aggregates, and differences in cerebral PSyn deposition profiles, we conclude that A53T_1_ and A53T_2_ PFFs propagate as two distinct conformational strains upon transmission to M83^+/-^ mice. A53T_2_ PFFs induced a more aggressive synucleinopathy characterized by faster disease kinetics and a higher extent of PSyn deposition, which is consistent to what we previously observed in M83^+/-^ mice inoculated with either S PFFs or MSA brain extract [47]. Given that the A53T_2_ PFF protofilament fold is also observed in α-syn fibrils extracted from MSA patient brain tissue (**Fig. 1i**), this provides a possible molecular explanation for why mice inoculated with A53T_2_ PFFs develop a more pronounced pathological phenotype.

**Figure 3.**
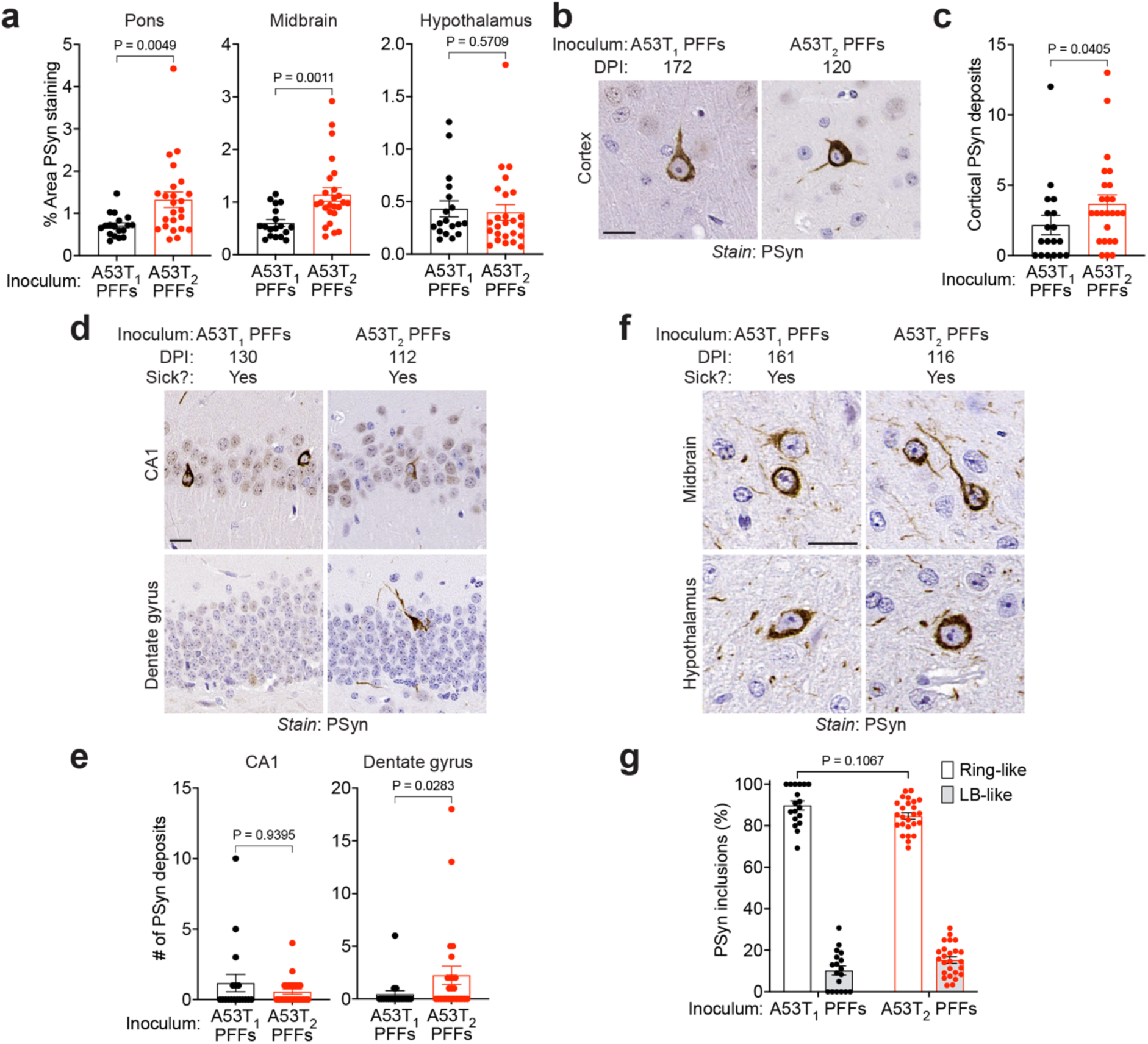
A53T_1_ and A53T_2_ PFFs induce distinct synucleinopathies in M83^+/-^ mice. **a**) Quantification of the area covered by PSyn staining in the indicated brain regions from symptomatic M83^+/-^ mice injected with A53T_1_ PFFs (n = 18) or A53T_2_ PFFs (n = 25). **b**) Representative images of cortical PSyn deposits in brain sections from M83^+/-^ mice inoculated with A53T_1_ or A53T_2_ PFFs. **c**) Quantification of the number of cortical PSyn deposits in brain sections from M83^+/-^ mice inoculated with A53T_1_ (n = 18) or A53T_2_ (n = 25) PFFs. **d**) Representative images of hippocampal PSyn deposits (CA1 and dentate gyrus regions) in brain sections from M83^+/-^ mice inoculated with A53T_1_ or A53T_2_ PFFs. **e**) Quantification of the number of PSyn deposits in the CA1 and dentate gyrus regions from M83^+/-^ mice inoculated with A53T_1_ (n = 18) or A53T_2_ (n = 25) PFFs. **f**) Representative images of ring-like PSyn deposits in the midbrain and hypothalamus from M83^+/-^ mice inoculated with A53T_1_ or A53T_2_ PFFs. **g**) Quantification of ring-like vs. Lewy body (LB)-like PSyn deposits in the midbrains of M83^+/-^ mice inoculated with A53T_1_ (n = 18) or A53T_2_ (n = 25) PFFs. In panels a, c, e, and g, the graphs display mean ± s.e.m. In panels a, c, and e, statistical significance was assessed by a Mann-Whitney test. In panel g, statistical significance was assessed by two-way ANOVA. In panels b, d, and f, the scale bar indicates 20 µm and applies to all images.

### Conformational variation of α-syn aggregates in spontaneously ill M83^+/+^ mice

While M83^+/-^ mice rarely develop a spontaneous synucleinopathy prior to 20 months of age, their homozygous counterparts (M83^+/+^) express higher amounts of A53T-mutant α-syn and thus exhibit spontaneous signs of motor dysfunction as early as 8 months of age [18]. Given that recombinant A53T-mutant α-syn could assemble into different aggregate structures, we wondered whether conformational variability among α-syn aggregates may also occur in the brains of M83^+/+^ mice. We aged 32 M83^+/+^ mice and allowed them to develop neurological symptoms. The onset of motor dysfunction in M83^+/+^ mice was highly variable, ranging from 211 to 596 days (**Fig. 4a**). To conformationally characterize the α-syn aggregates present in the brains of spontaneously ill M83^+/+^ mice, we conducted protease digestion assays with TL and PK. Remarkably, based on the patterns of protease-resistant α-syn species observed in brain extracts, we found that spontaneously ill M83^+/+^ mice could be classified into three disease subtypes, which we termed M83^+/+^_X_, M83^+/+^_Y_, and M83^+/+^_Z_ (**Fig. 4b**). M83^+/+^_X_ was characterized by a single TL-resistant α-syn band at ∼15 kDa, and two equally strong PK-resistant α-syn bands at ∼11 and ∼10 kDa. M83^+/+^_Y_ was characterized by two TL-resistant α-syn bands at ∼16 and ∼15 kDa, and one dominant PK-resistant α-syn band at ∼10 kDa. Like the M83^+/+^_Y_ subtype, M83^+/+^_Z_ was characterized by two TL-resistant α-syn bands at ∼16 and ∼15 kDa. However, M83^+/+^_Z_ possessed a dominant PK-resistant α-syn band at ∼11 kDa (instead of ∼10 kDa). Of the three subtypes, M83^+/+^_Y_ was by far the most common followed by M83^+/+^_X_ and then M83^+/+^_Z_ (**Fig. 4c**). We wondered whether the occurrence of different disease subtypes was linked to the sex or age of disease onset in the M83^+/+^ mice. However, both male and female mice developed all three synucleinopathy subtypes, and the ages of disease onset were not statistically different across the three subtypes (**Fig. 4d**).

**Figure 4.**
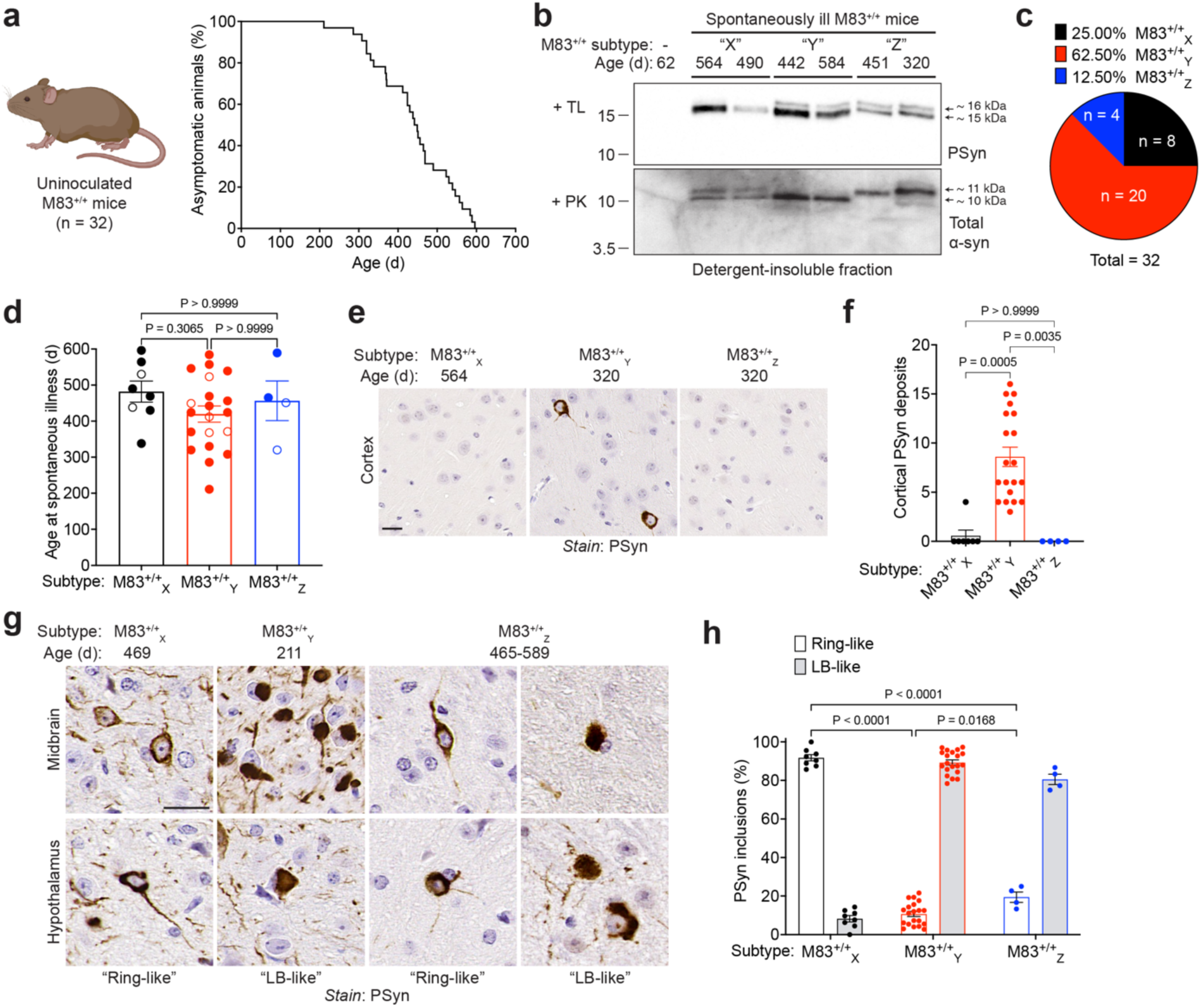
Spontaneously ill M83^+/+^ mice can be classified into three disease subtypes. **a**) Kaplan-Meier survival curve for M83^+/+^ mice (n = 32). **b**) Immunoblots of detergent-insoluble TL-resistant PSyn species and PK-resistant total α-syn species in brain extracts from spontaneously ill M83^+/+^ mice at the indicated ages. Based on the patterns of TL- and PK-resistant species, each mouse was assigned as either subtype X, Y, or Z. Two representative mice for each subtype are shown. Brain extract from a young asymptomatic M83^+/+^ mouse is included as a negative control. Molecular weight markers indicate kDa. **c**) Relative frequencies of M83^+/+^_X_, M83^+/+^_Y_, and M83^+/+^_Z_ subtypes across the 32 observed M83^+/+^ mice. **d**) Age of spontaneous motor dysfunction onset in M83^+/+^ mice classified as M83^+/+^_X_ (n = 8), M83^+/+^_Y_ (n = 20), or M83^+/+^_Z_ (n = 4). Male mice are indicated by open circles and female mice by closed circles. **e**) Representative images of cortical PSyn deposits in brain sections from spontaneously sick mice exhibiting the M83^+/+^_X_, M83^+/+^_Y_, or M83^+/+^_Z_ subtype. **f**) Quantification of the number of cortical PSyn deposits in brain sections from spontaneously sick mice displaying the M83^+/+^_X_ (n = 8), M83^+/+^_Y_ (n = 20), or M83^+/+^_Z_ (n = 4) subtype. **g**) Representative images of ring-like and Lewy body (LB)-like PSyn deposits in the midbrain and hypothalamus from spontaneously ill M83^+/+^ mice. **h**) Quantification of ring-like vs. LB-like PSyn deposits in the midbrains of spontaneously ill M83^+/+^ mice exhibiting the M83^+/+^_X_ (n = 8), M83^+/+^_Y_ (n = 20), or M83^+/+^_Z_ (n = 4) subtype. All graphs display mean ± s.e.m. In panels d and f, statistical significance was assessed by a Kruskal-Wallis test followed by Dunn’s multiple comparisons test. In panel h, statistical significance was assessed by two-way ANOVA. In panels e and g, the scale bar indicates 20 µm and applies to all images.

Levels of detergent-insoluble PSyn in brain extracts were indistinguishable across the three M83^+/+^ disease subtypes (**Supplemental Fig. 5a**). Similarly, levels of PSyn deposition tended to be higher in the pons, midbrain, and hypothalamus of M83^+/+^_Y_ mice, but there was considerable animal-to-animal variability (**Supplemental Fig. 5b**). However, a defining feature of M83^+/+^_Y_ mice was the occurrence of PSyn deposits in cortical neurons, while the brains of M83^+/+^_X_ and M83^+/+^_Z_ mice had mostly none (**Fig. 4e, f**). In addition, striking differences in the morphology of neuronal PSyn deposits in the midbrain and hypothalamus were observed across the three M83^+/+^ disease subtypes. M83^+/+^_X_ mice had predominantly ring-like aggregates, M83^+/+^_Y_ mice had predominantly dense LB-like aggregates, and while M83^+/+^_Z_ mice also had more LB-like than ring-like aggregates, a greater proportion of ring-like aggregates were present than in M83^+/+^_Y_ mice (**Fig. 4g, h**). Therefore, aged M83^+/+^ mice spontaneously develop one of three distinct disease subtypes characterized by differences in the banding patterns of protease-resistant α-syn species, targeting of the cerebral cortex, and the morphology of cerebral PSyn deposits.

### M83^+/+^ subtypes transmit as distinct α-syn strains in M83^+/-^ mice

To determine if the three M83^+/+^ subtypes propagate as distinct strains, we intracerebrally inoculated brain homogenates from three spontaneously ill M83^+/+^ mice per subtype into M83^+/-^ mice, with brain homogenate from a young asymptomatic M83^+/+^ mouse serving as a negative control (**Fig. 5a**). Whereas all control-inoculated mice remained asymptomatic up to 540 DPI, 69/71 M83^+/-^ mice inoculated with spontaneously sick M83^+/+^ brain extract developed progressive motor impairment (**Supplemental Table 2**). M83^+/-^ mice injected with M83^+/+^_X_ brain extract developed disease significantly faster than mice injected with either M83^+/+^_Y_ or M83^+/+^_Z_ brain extract (**Fig. 5b**). Moreover, M83^+/-^ mice injected with M83^+/+^_X_ had significantly higher overall amounts of detergent-insoluble PSyn species in their brains than mice injected with either M83^+/+^_Y_ or M83^+/+^_Z_ (**Supplemental Fig. 6a**). We next compared the banding patterns of protease-resistant α-syn species in brain extracts from the different groups of inoculated M83^+/-^ mice following TL or PK digestion. Following inoculation into M83^+/-^ mice, the defining biochemical characteristics of each M83^+/+^ subtype were maintained (**Fig. 5c**). For example, like M83^+/+^_X_ mice, M83^+/+^_X_-inoculated M83^+/-^ mice had one TL-resistant band at ∼15 kDa and two dominant PK-resistant bands at ∼11 kDa and ∼10 kDa. Similarly, mimicking the differences between M83^+/+^_Y_ and M83^+/+^_Z_ mice, M83^+/+^_Y_-inoculated M83^+/-^ mice had a dominant PK-resistant band at ∼10 kDa whereas M83^+/+^_Z_-inoculated M83^+/-^ mice had a dominant PK-resistant band at ∼11 kDa.

**Figure 5.**
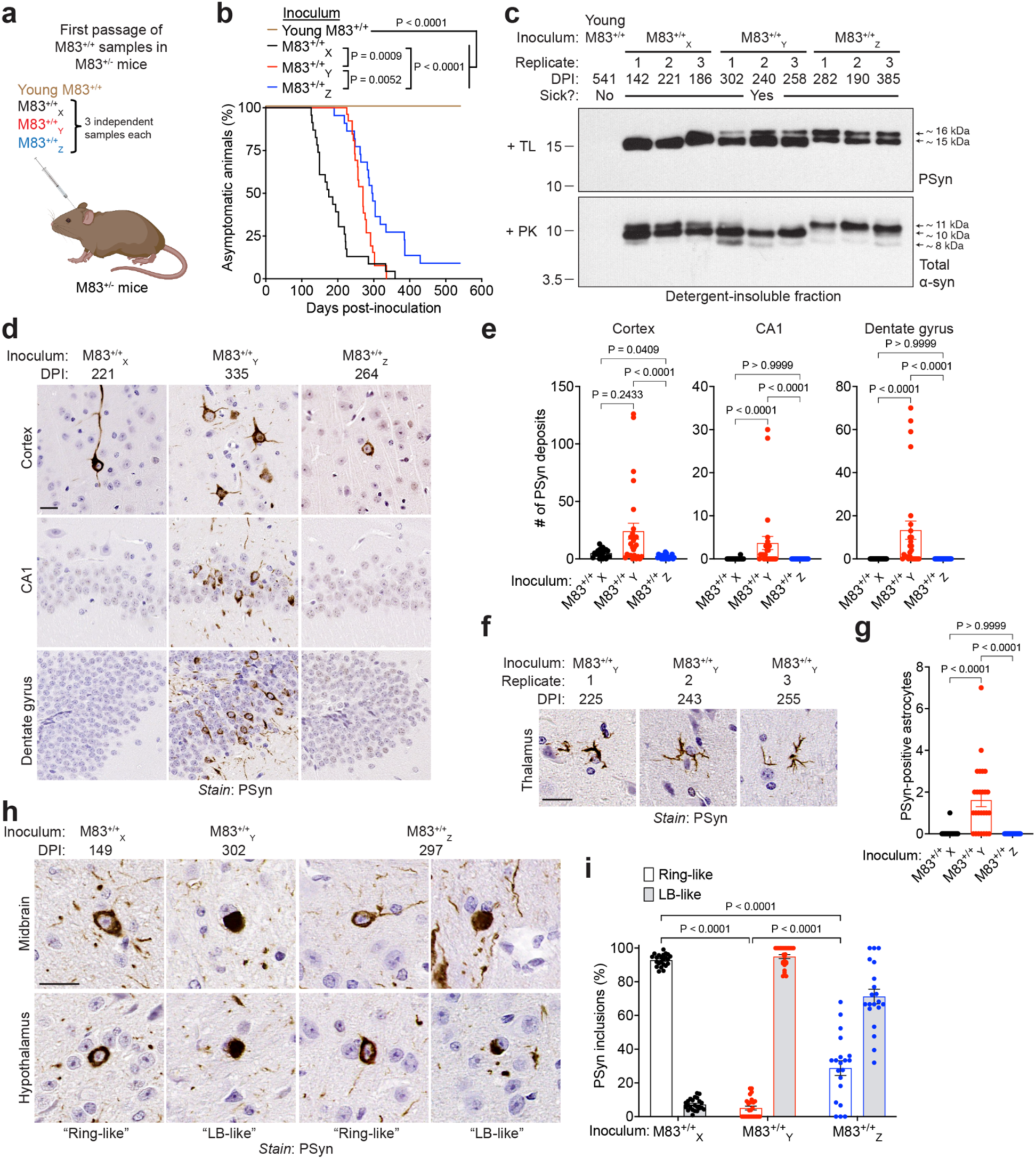
Propagation of M83^+/+^ subtypes in M83^+/-^ mice. **a**) Schematic of intracerebral inoculation experiments in M83^+/-^ mice. **b**) Kaplan-Meier survival curves for M83^+/-^ mice inoculated with young M83^+/+^ brain extract (n = 8) or with symptomatic M83^+/+^ brain extract from mice exhibiting the M83^+/+^_X_ (n = 23), M83^+/+^_Y_ (n = 26), or M83^+/+^_Z_ (n = 22) subtype. For the symptomatic M83^+/+^ inoculations, data from the three independent experiments for each subtype is pooled. Statistical significance was determined using the Log-rank test. **c**) Immunoblots of detergent-insoluble TL-resistant PSyn species and PK-resistant total α-syn species in brain extracts from symptomatic inoculated M83^+/-^ mice at the indicated DPI. One representative mouse from each of 3 replicate experiments for each M83^+/+^ subtype is shown. Brain extract from an asymptomatic M83^+/-^ mouse inoculated with a young M83^+/+^ sample is included as a negative control. Molecular weight markers indicate kDa. **d**) Representative images of cortical and hippocampal (CA1 and dentate gyrus regions) PSyn deposits in brain sections from symptomatic M83^+/-^ mice inoculated with either of the 3 different M83^+/+^ subtypes. **e**) Quantification of the number of cortical, CA1, and dentate gyrus PSyn deposits in brain sections from symptomatic M83^+/-^ mice inoculated with either the M83^+/+^_X_ (n = 23), M83^+/+^_Y_ (n = 26), or M83^+/+^_Z_ (n = 20) subtype. **f**) Representative images of astrocytic PSyn deposits in the thalamus of symptomatic M83^+/-^ mice inoculated with either of 3 replicates of M83^+/+^_Y_ brain extract. **g**) Quantification of PSyn-positive astrocytes in brain sections from symptomatic M83^+/-^ mice inoculated with either the M83^+/+^_X_ (n = 23), M83^+/+^_Y_ (n = 26), or M83^+/+^_Z_ (n = 20) subtype. **h**) Representative images of ring-like and Lewy body (LB)-like PSyn deposits in the midbrain and hypothalamus from symptomatic M83^+/-^ mice inoculated with either of the 3 M83^+/+^ subtypes. **i**) Quantification of ring-like vs. LB-like PSyn deposits in the midbrains of symptomatic M83^+/-^ mice inoculated with either the M83^+/+^_X_ (n = 23), M83^+/+^_Y_ (n = 26), or M83^+/+^_Z_ (n = 20) disease subtype. All graphs display mean ± s.e.m. In panels e and g, statistical significance was assessed by a Kruskal-Wallis test followed by Dunn’s multiple comparisons test. In panel i, statistical significance was assessed by two-way ANOVA. In panels d, f, and h, the scale bar indicates 20 µm and applies to all images.

As expected, all groups of M83^+/-^ mice inoculated with brain extract from spontaneously sick M83^+/+^ mice exhibited significantly higher levels of cerebral PSyn deposition than M83^+/-^ mice injected with brain extract from a young asymptomatic M83^+/+^ mouse (**Supplemental Fig. 6b**). Consistent with the increased amounts of detergent-insoluble PSyn species observed in M83^+/+^_X_-injected mice, significantly higher levels of PSyn pathology were present in the midbrain of M83^+/+^_X_-injected mice compared to either M83^+/+^_Y_-or M83^+/+^_Z_-injected animals (**Supplemental Fig. 6c**). However, M83^+/+^_Y_-injected M83^+/-^ mice had significantly more PSyn deposition than mice injected with either M83^+/+^_X_ or M83^+/+^_Z_ in the cerebral cortex, CA1, and dentate gyrus (**Fig. 5d, e**). The brains of M83^+/-^ mice inoculated with M83^+/+^_Y_ also uniquely exhibited glial PSyn deposition in the thalamus (**Fig. 5f, g**). Thus, despite overall lower levels of cerebral PSyn accumulation, M83^+/+^_Y_-injected M83^+/-^ mice exhibited selective cellular and brain region targeting of PSyn aggregates. Finally, consistent with the conserved patterns of protease-resistant α-syn species, the morphology of neuronal PSyn inclusions in the midbrain and hypothalamus of M83^+/-^ mice matched that of the M83^+/+^ subtype with which they were injected (**Fig. 5h, i**). For instance, M83^+/+^_X_-inoculated mice had mostly ring-like deposits, M83^+/+^_Y_-inoculated mice had mostly LB-like deposits, and M83^+/+^_Z_-inoculated mice possessed a mixture of both. Overall, these findings suggest that the strain-specific structural characteristics of α-syn aggregates from the three M83^+/+^ subtypes were faithfully preserved upon propagation in M83^+/-^ mice, likely due to templated conformational conversion.

### Serial propagation of M83^+/+^ subtypes in M83^+/-^ mice

To investigate whether the unique characteristics of the individual M83^+/+^ subtypes are stable, as would be expected if they constitute distinct α-syn strains, we performed second passage experiments in M83^+/-^ mice. One brain homogenate sample from the initial passage of each M83^+/+^ subtype into M83^+/-^ mice was selected for inoculation into a second set of M83^+/-^ mice (**Fig. 6a**). These inocula were termed M83^+/-^_X_, M83^+/-^_Y_, and M83^+/-^_Z_. As in the initial passage, M83^+/-^ mice inoculated with M83^+/-^_X_ developed neurological illness significantly more rapidly than mice inoculated with either M83^+/-^_Y_ or M83^+/-^_Z_ (**Fig. 6b**). All the inoculated M83^+/-^ second passage mice developed neurological disease (**Supplemental Table 3**). As in the first passage, higher levels of detergent-insoluble PSyn were found in brain homogenates from mice inoculated with M83^+/-^_X_ than mice inoculated with M83^+/-^_Z_ (**Supplemental Fig. 7a**). Next, we performed protease digestion assays to determine whether strain-specific banding pattern differences were maintained upon second passage. Indeed, for each M83^+/+^ subtype, the banding patterns of protease-resistant α-syn species in the brains of second passage M83^+/-^ mice were identical to those from the first passage mice as well as the original M83^+/+^ samples (**Fig. 6c**).

**Figure 6.**
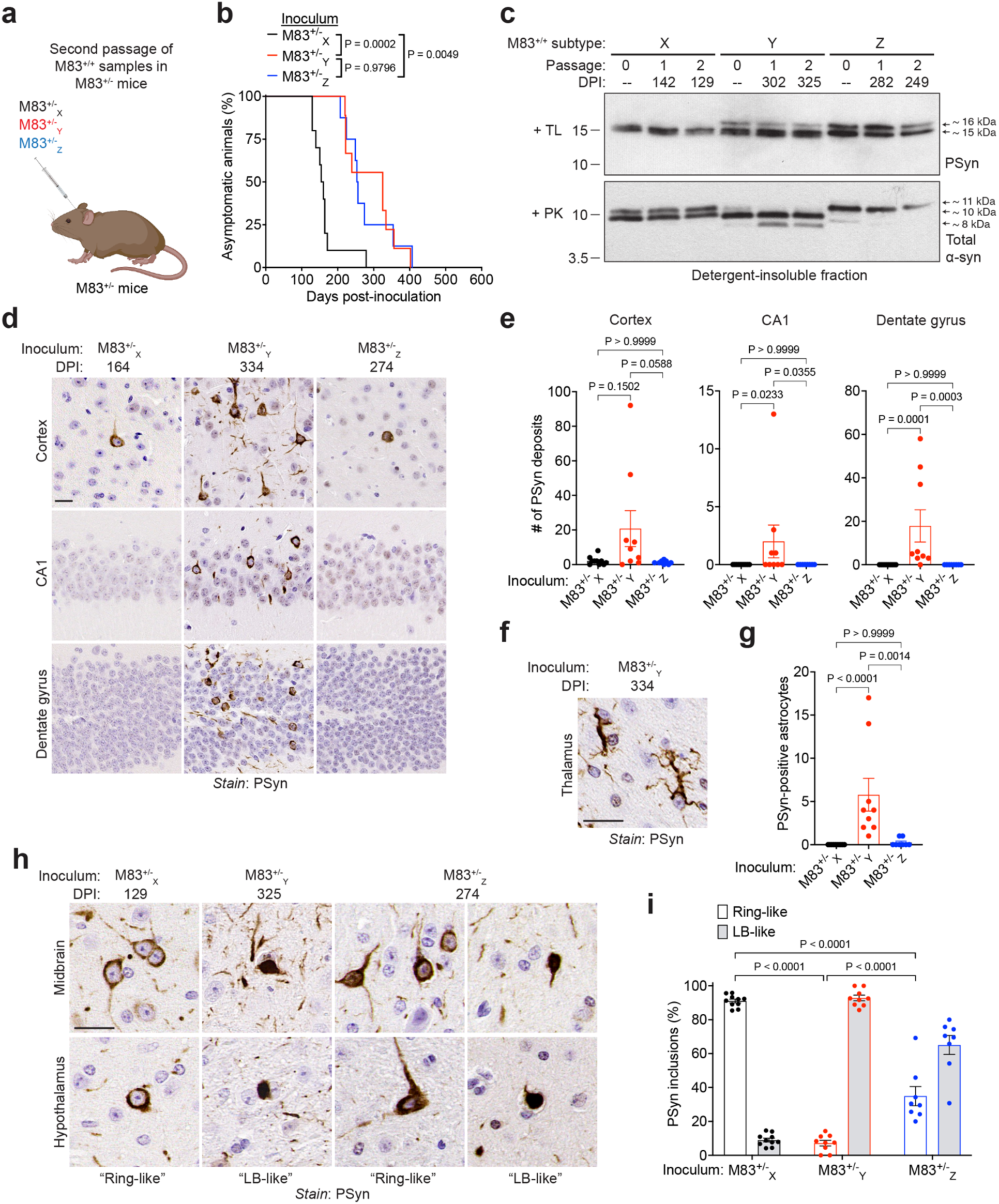
Second passage of M83^+/+^ subtypes in M83^+/-^ mice. **a**) Schematic of second passage experiments in M83^+/-^ mice. Mice were inoculated with brain extract from symptomatic M83^+/-^ mice that were previously inoculated with either the X (M83^+/-^_X_), Y (M83^+/-^_Y_), or Z (M83^+/-^_Z_) M83^+/+^ subtype. **b**) Kaplan-Meier survival curves for M83^+/-^ mice inoculated with M83^+/-^_X_ (n = 10), M83^+/-^_Y_ (n = 9), or M83^+/-^_Z_ (n = 8) brain extract. Statistical significance was determined using the Log-rank test. **c**) Immunoblots of detergent-insoluble TL-resistant PSyn species and PK-resistant total α-syn species in brain extracts from spontaneously ill M83^+/+^ mice (“passage 0”) or symptomatic M83^+/-^ mice at the indicated DPI following the first or second passage of the M83^+/+^_X_, M83^+/+^_Y_, or M83^+/+^_Z_ subtypes. Molecular weight markers indicate kDa. **d**) Representative images of cortical and hippocampal (CA1 and dentate gyrus regions) PSyn deposits in brain sections from symptomatic M83^+/-^ mice inoculated with the indicated samples. **e**) Quantification of the number of cortical, CA1, and dentate gyrus PSyn deposits in brain sections from symptomatic M83^+/-^ mice inoculated with either M83^+/-^_X_ (n = 10), M83^+/-^_Y_ (n = 9), or M83^+/-^_Z_ (n = 8) brain homogenate. **f**) Representative image of astrocytic PSyn deposits in the thalamus of a symptomatic M83^+/-^ mouse inoculated with M83^+/-^_Y_ brain extract. **g**) Quantification of PSyn-positive astrocytes in brain sections from symptomatic M83^+/-^ mice inoculated with either M83^+/-^_X_ (n = 10), M83^+/-^_Y_ (n = 9), or M83^+/-^_Z_ (n = 8) brain homogenate. **h**) Representative images of ring-like (black arrows) and Lewy body (LB)-like (red arrows) PSyn deposits in the midbrain and hypothalamus from symptomatic M83^+/-^ mice inoculated with the indicated samples. **i**) Quantification of ring-like vs. LB-like PSyn deposits in the midbrains of symptomatic M83^+/-^ mice inoculated with either M83^+/-^_X_ (n = 10), M83^+/-^_Y_ (n = 9), or M83^+/-^_Z_ (n = 8) brain homogenate. All graphs display mean ± s.e.m. In panels e and g, statistical significance was assessed by a Kruskal-Wallis test followed by Dunn’s multiple comparisons test. In panel i, statistical significance was assessed by two-way ANOVA. In panels d, f, and h, the scale bar indicates 20 µm and applies to all images.

Similar to what was observed in the initial passage of M83^+/+^ subtypes in M83^+/-^ mice, mice injected with M83^+/-^_Y_ also developed the most PSyn inclusions in the cerebral cortex, CA1, and dentate gyrus compared to mice injected with either M83^+/-^_X_ or M83^+/-^_Z_ (**Fig. 6d, e**), despite the fact that comparable levels of PSyn deposition were observed in other brain regions (**Supplemental Fig. 7b**). M83^+/-^_Y_-injected mice also uniquely exhibited glial PSyn inclusions in the thalamus (**Fig. 6f, g**). Furthermore, the M83^+/+^ subtype-specific morphological differences in PSyn inclusions within midbrain and hypothalamus neurons were conserved upon second passage: M83^+/-^_X_-inoculated mice had predominantly ring-like aggregates, M83^+/-^_Y_-inoculated mice had predominantly LB-like aggregates, and M83^+/-^_Z_-inoculated mice had a more even distribution of both aggregate types (**Fig. 6h, i**). Therefore, we can conclude that the three M83^+/+^ disease subtypes consist of three different α-syn conformational strains that arise naturally, and that α-syn strain-specific properties are stable and faithfully maintained during serial passaging. What we referred to as the “M83” strain in our previous study appears to be identical to the M83^+/+^_Y_ subtype [47].

### Recombinant human WT α-syn assembles into at least 6 distinct polymorphs

After characterizing strains made from A53T-mutant α-syn *in vitro* and *in vivo*, we next analyzed potential conformational variability in PFFs made from WT α-syn, as sporadic PD and MSA cases are characterized by the presence of WT α-syn aggregates in the brain. To generate WT α-syn PFFs, we polymerized recombinant human WT α-syn using the same salt-containing buffer and shaking conditions we used to generate A53T PFFs (**Fig. 7a**). We generated 35 independent WT PFF preparations and then characterized their conformational properties by PK digestion. Based on the patterns of PK-resistant α-syn fragments observed, we found that individual preparations of WT PFFs could be classified into 6 different groups (**Fig. 7b**). We named these polymorphs WT_1_ to WT_6_; multiple instances of the WT_1_, WT_3_, WT_4_, and WT_5_ PFF polymorphs were observed while the WT_2_ and WT_6_ polymorphs were only obtained a single time (**Fig. 7c**). Multiple PFF polymorphs were obtained from each of four independent batches of recombinant WT human α-syn (**Supplemental Fig. 8a**). Using the curcumin dye-binding assay, we found that WT_6_ PFFs had a completely different fluorescence emission spectrum compared to the other polymorphs, suggestive of a structural difference (**Supplemental Fig. 8b**). Abundant fibrils were observed in WT PFF preparations by negative stain electron microscopy (**Supplemental Fig. 8c**).

**Figure 7.**
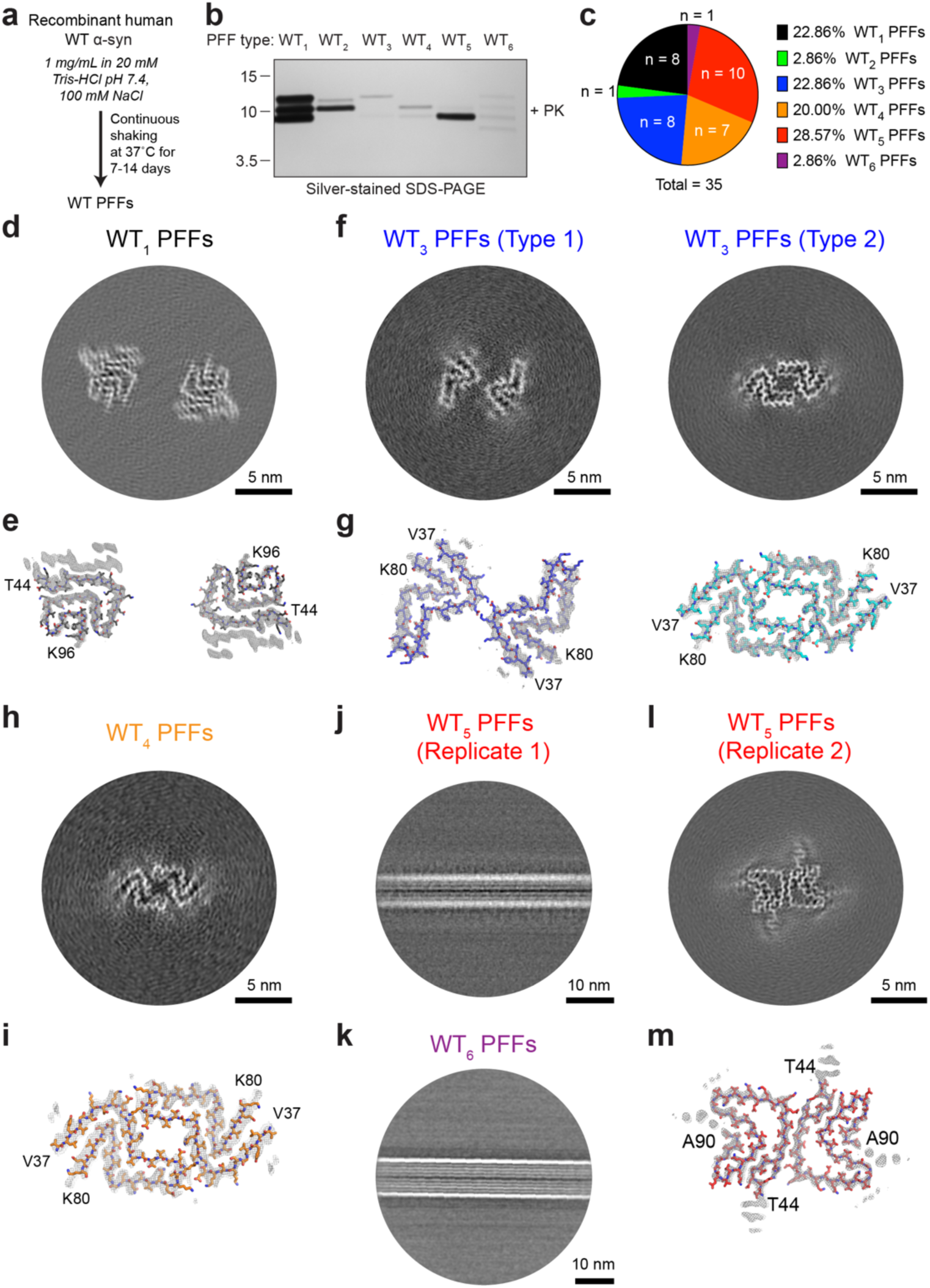
Wild-type recombinant α-syn forms many distinct PFF polymorphs. **a**) Schematic for formation of WT PFFs. **b**) Following PK digestion of WT PFFs and analysis by SDS-PAGE, differences in the pattern of PK-resistant insoluble α-syn fragments reveals six distinct PFF polymorphs (WT_1_ to WT_6_). Molecular weight markers indicate kDa. **c**) Relative frequencies of WT PFF polymorphs across 35 independent PFF preparations. **d, f, h, l**) Cross-sections of the cryo-EM density maps showing WT_1_ PFFs (d), WT_3_ Type 1 and Type 2 PFFs (f), WT_4_ PFFs (h), and WT_5_ PFFs (l). Scale bars indicate 5 nm. **e, g, i, m**) Overlay of sharpened high-resolution cryo-EM density maps (gray mesh representation) and the atomic models for the various WT PFF polymorphs. **j, k**) Representative 2D class images for a distinct preparation of WT_5_ PFFs as well as for WT_6_ PFFs, for both of which only straight, non-twisted fibrils were observed.

We next used cryo-EM to investigate the molecular architecture of α-syn fibrils within the WT_1_, WT_3_, WT_4_, WT_5_, and WT_6_ polymorphs. Due to insufficient sample availability, we were unable to perform further structural characterization of the single preparation of WT_2_ PFFs obtained. As with the A53T PFFs, all WT α-syn PFF polymorphs consisted of two intertwined protofilaments. The filament structure of WT_1_ PFFs was atypical in that a large inter-protofilament distance was observed, suggestive of an absence of fibril-stabilizing inter-protofilament interactions (**Fig. 7d, e**). For the WT_3_ polymorph, two different fibril types were observed. Both possessed the same core α-syn structure but differed in the arrangement of protofilaments and the inter-protofilament contact sites (**Fig. 7f, g**). WT_4_ PFFs adopted a similar molecular structure to that of WT_3_ PFFs, but only a single arrangement of the two protofilaments was found (**Fig. 7h, i**). As structural determination of protein fibrils by cryo-EM requires that fibrils exhibit a twist [52], we were initially unable to determine the structures of WT_5_ PFFs as well as the single preparation of WT_6_ PFFs obtained due to the exclusive presence of straight fibrils (**Fig. 7j, k**). Thus, we attempted cryo-EM analysis using a second replicate of WT_5_ PFFs, which yielded twisted fibrils, allowing us to obtain a molecular structure. The molecular structure of this replicate of WT_5_ PFFs was distinct from that of both WT_1_ and WT_3_ PFFs (**Fig. 7l, m**). We therefore conclude that diverse fibrillar structures emerge among individual preparations of WT α-syn PFFs formed under identical conditions.

To determine if individual WT PFF polymorphs initiate distinct synucleinopathies, we intracerebrally inoculated groups of M83^+/-^ mice with one or two preparations of each PFF type (**Fig. 8a**). All samples that were investigated by cryo-EM were inoculated. Despite the sequence mismatch between WT α-syn in the PFFs and A53T-mutant α-syn expressed by M83^+/-^ mice, experiments were performed in this line to enable direct comparison with the transmissions involving A53T PFF polymorphs and M83^+/+^ disease subtypes. Based on the disease kinetics observed, we found that the WT PFF polymorphs could be classified into three distinct clusters: WT_1_, WT_2_, and WT_4_ PFFs caused rapid disease onset with incubation periods of ∼4 months, WT_5_ PFFs caused a more slowly progressive neurological disease with an incubation period of ∼7-8 months, whereas WT_3_ and WT_6_ PFFs resulted in inefficient disease transmission with most mice remaining asymptomatic for up to 18 months post-inoculation (**Fig. 8b, Supplemental Table 4**). None of the M83^+/-^ mice inoculated with monomeric WT α-syn developed neurological illness. Consistent with their similar disease incubation periods, the TL and PK digestion profiles of α-syn aggregates in brain extracts from mice injected with either WT_1_, WT_2_, or WT_4_ PFFs were identical (**Supplemental Fig. 9a**). Moreover, the histological patterns of PSyn deposition in the brains of M83^+/-^ mice inoculated with WT_1_, WT_2_, or WT_4_ PFFs were comparable (**Supplemental Fig. 9b, c**). Thus, despite exhibiting distinct profiles of PK-resistant α-syn fragments, WT_1_, WT_2_, and WT_4_ PFFs produced a highly similar synucleinopathy upon inoculation into M83^+/-^ mice.

**Figure 8.**
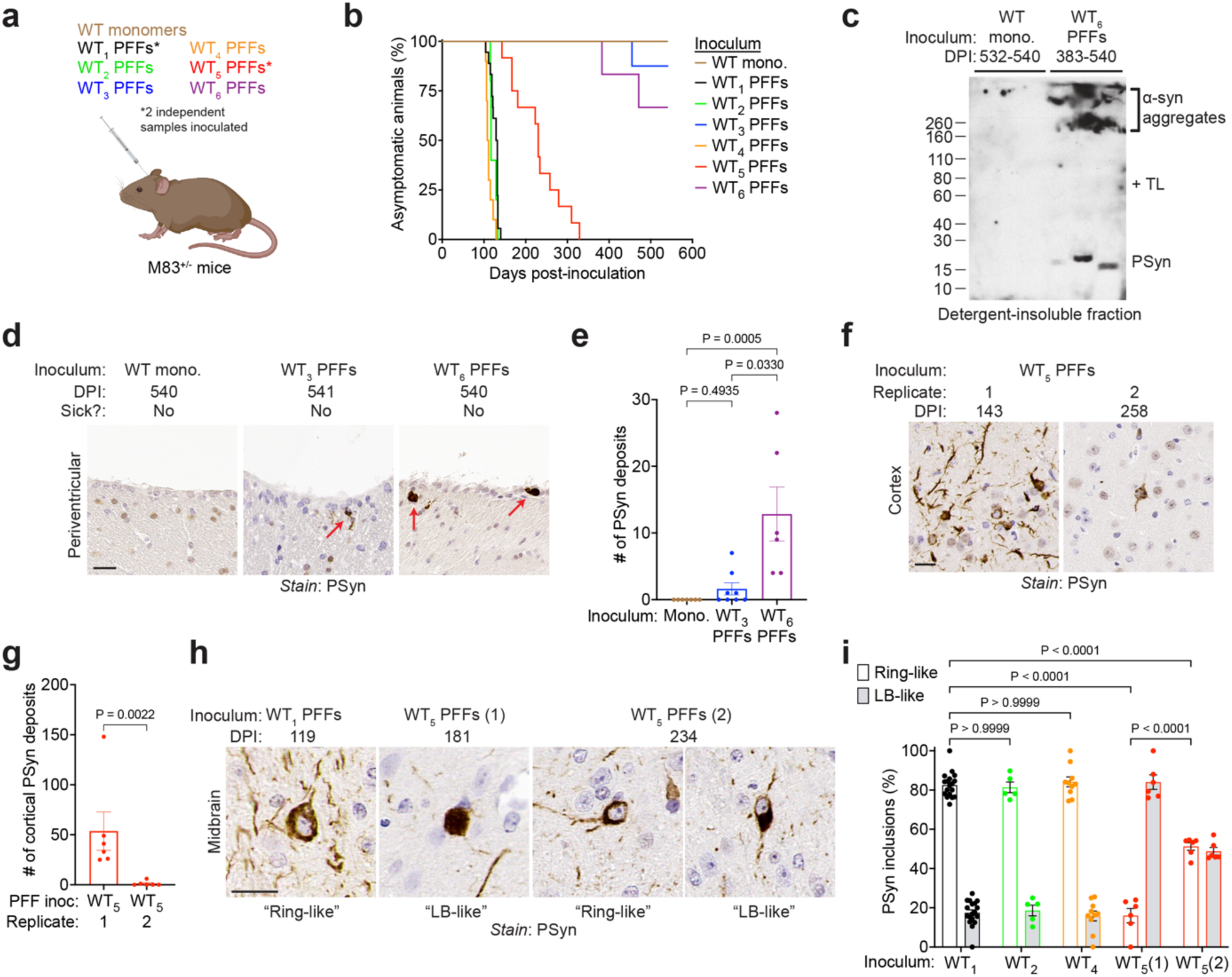
Wild-type α-syn PFF polymorphs induce diverse synucleinopathies in M83^+/-^ mice. **a**) Schematic of intracerebral inoculation experiments in M83^+/-^ mice. **b**) Kaplan-Meier survival curves for M83^+/-^ mice inoculated with monomeric (mono.) WT α-syn (n = 7) or with WT_1_ (n = 18), WT_2_ (n = 5), WT_3_ (n = 8), WT_4_ (n = 10), WT_5_ (n = 12), or WT_6_ (n = 6) PFFs. For the WT_1_ and WT_5_ PFF inoculations, data from two replicate experiments for each polymorph is pooled. **c**) Immunoblot of detergent-insoluble PSyn species in TL-digested brain extracts from M83^+/-^ mice inoculated with either monomeric α-syn or WT_5_ PFFs. Samples from three independent mice per inoculum are shown. Molecular weight markers indicate kDa. **d**) Representative images of periventricular PSyn deposits (red arrows) in brain sections from M83^+/-^ mice inoculated with WT_3_ or WT_6_ PFFs. A brain section from a mouse inoculated with monomeric α-syn is shown as a control. **e**) Quantification of the number of periventriclar PSyn deposits in brain sections from M83^+/-^ mice inoculated with monomeric α-syn (n = 7), WT_3_ PFFs (n = 8), or WT_6_ PFFs (n = 6). Statistical significance was assessed using a Kruskal-Wallis test with Dunn’s multiple comparisons test. **f**) Representative images of cortical PSyn deposits in brain sections from symptomatic M83^+/-^ mice inoculated with either of two replicates of WT_5_ PFFs. **g**) Quantification of the number of cortical PSyn deposits from M83^+/-^ mice inoculated with two independent preparations of WT_5_ PFFs (n = 6 each). **h**) Representative images of ring-like and Lewy body (LB)-like PSyn deposits in the midbrain of symptomatic M83^+/-^ mice inoculated with the indicated samples. **i**) Quantification of ring-like vs. LB-like PSyn deposits in the midbrains of symptomatic M83^+/-^ mice inoculated with either WT_1_ (n = 18), WT_2_ (n = 5), or WT_4_ (n = 10) PFFs, or with either of two different replicates of WT_5_ PFFs (n = 6 each). In panels e, g, and i, the graphs display mean ± s.e.m. In panels d, f, and h, the scale bar indicates 20 µm and applies to all images.

While most M83^+/-^ mice inoculated with WT_6_ PFFs did not develop overt neurological symptoms within the experimental timeframe, we found low amounts of detergent-insoluble and TL-resistant PSyn aggregates in brain extracts from mice injected with WT_6_ PFFs but not in extracts from mice at 18 months post-inoculation with monomeric WT α-syn (**Fig. 8c**). While some of the TL-resistant PSyn species migrated near their predicted molecular weight (∼15 kDa), most of the signal remained near the top of the gel, suggestive of the presence of SDS-resistant α-syn aggregates in the brains of mice injected with WT_6_ PFFs. Consistent with this observation, half of the WT_3_ PFF-injected mice and all the WT_6_ PFF-injected animals developed sparse PSyn aggregates near the interface between the brain parenchyma and the ventricular system (**Fig. 8d, e**). All PSyn deposits in mice injected with WT_6_ PFFs were large and dense LB-like inclusions; such deposits were absent in the brains of M83^+/-^ mice inoculated with monomeric WT α-syn. Thus, WT_6_ PFFs induced a very different synucleinopathy in the brains of M83^+/-^ mice than that induced by any of the other WT or A53T-mutant α-syn PFF polymorphs.

Despite both WT_5_ PFF preparations used for inoculation experiments exhibiting an identical pattern of PK-resistant α-syn fragments and initiating a slower disease upon inoculation into M83^+/-^ mice (**Supplemental Fig. 10a, Supplemental Table 4**), we found inter-replicate differences in the synucleinopathies induced. For instance, after propagation of WT_5_ PFFs in M83^+/-^ mice, the induced α-syn aggregates exhibited replicate-specific differences in their PK and TL digestion patterns (**Supplemental Fig. 10b, c**). Specifically, we found that the ratio of 16 kDa/15 kDa TL-resistant fragments was significantly lower and the ratio of 8 kDa/10 kDa PK-resistant fragments significantly higher in brain extracts from mice inoculated with Replicate 1 of WT_5_ PFFs than in brain extracts from mice inoculated with Replicate 2. Further differences were observed upon histopathological analysis. Replicate 1 of WT_5_ PFFs induced abundant neuronal PSyn aggregates in the cortex while Replicate 2 did not (**Fig. 8f, g**). The morphology of neuronal PSyn deposits in the midbrain was also distinct: Replicate 1 produced predominantly LB-like aggregates, while Replicate 2 produced roughly equal proportions of ring-like and LB-like inclusions (**Fig. 8h, i**). In contrast, the brains of M83^+/-^ mice inoculated with either WT_1_, WT_2_, or WT_4_ PFFs all displayed predominantly ring-like PSyn deposits. Therefore, WT α-syn PFFs can induce at least three different synucleinopathies in M83^+/-^ mice, and distinct diseases can even emerge when replicates of a single seemingly uniform PFF type are propagated *in vivo*.

### A53T PFFs, WT PFFs, and M83^+/+^ aggregates form overlapping α-syn strains

During the characterization of groups of inoculated M83^+/-^ mice by TL and PK digestion, we noticed that mice from different experiments shared similar banding patterns of protease-resistant α-syn species and histopathological features. To look for overlapping strain types, we collectively analyzed brain homogenates from mice inoculated with either A53T_1_ PFFs, A53T_2_ PFFs, WT_1_ PFFs, WT_5_ PFFs (Replicate 1), M83^+/+^_X_, M83^+/+^_Y,_ or M83^+/+^_Z_ using TL and PK digestion (**Supplemental Fig. 11**). Based on the patterns of protease-resistant α-syn fragments obtained, three distinct strain types could be identified, and each of these strain types was associated with a specific set of disease attributes (**Supplemental Table 5**). Strain type 1 could be generated by inoculation with either A53T PFFs, WT PFFs, or M83^+/+^ brain extract whereas strain type 2 could only be achieved through inoculation with WT PFFs or M83^+/+^ homogenate. Strain type 1 appears to be very similar, if not identical to the strain type present in M83^+/-^ mice inoculated with either S PFFs or MSA brain extract, whereas strain type 2 appears to be the same as in M83^+/-^ mice inoculated with NS PFFs [47]. Strain type 3, which we have not previously observed, was generated either by inoculation with A53T PFFs or with M83^+/+^ brain extract. A completely different synucleinopathy was observed in M83^+/-^ mice inoculated with WT_6_ PFFs, which we have designated as strain type 4. Although the low levels of α-syn aggregates present in the brains of WT_6_ PFF-injected mice prevented us from performing strain typing by PK and TL digestion, the strain type 4 phenotype is highly similar to that observed in M83^+/-^ mice inoculated with brain extract from PD patients [53, 54]. Thus, M83^+/-^ mice are capable of propagating at least four distinct strains of α-syn aggregates.

## Discussion

In this study, we demonstrate that a consistent molecular environment can give rise to structurally diverse α-syn aggregates that induce distinct synucleinopathies upon propagation in a mouse model. For both WT and A53T-mutant recombinant α-syn, we observed structural diversity in PFFs generated under identical conditions across several batches of purified protein, and there was no way of predicting the specific polymorph that would occur in a single PFF preparation. Similarly, using inbred transgenic mice to minimize inter-animal genetic differences, we found that three distinct synucleinopathy subtypes emerge spontaneously in M83^+/+^ mice, and specific disease subtypes were not linked to defined mouse attributes such as age or sex. Thus, given the seemingly random nature of α-syn strain generation both *in vitro* and *in vivo*, we conclude that stochastic misfolding of α-syn into one of several possible aggregate structures is a major driver of disease heterogeneity in the synucleinopathies. As we did not observe any evidence of mixtures of different α-syn strains among the three M83^+/+^ subtypes, either in spontaneously sick animals or upon serial propagation in M83^+/-^ mice, it is conceivable that a single initial stochastic misfolding event defines the entire downstream disease course. This could result from different α-syn strains actively interfering with each other’s propagation, as can occur with prion strains [55], or because spontaneous formation of a stable, self-propagating α-syn aggregate seed is a rare event that precedes rapid seed-induced templating and cell-to-cell spreading of α-syn aggregates in the brain [56].

Potential stochastic formation of structurally diverse protein aggregates has also been observed in other studies. In one report, an experimental condition previously shown to generate ThT-positive α-syn fibrils also gave rise to ThT-negative polymorphs that exhibited different seeding properties in cultured cells and mice [49]. In another study, α-syn aggregation intermediates in the same tube could be separated into two distinct fractions by their solubility, and they proceeded to develop into structurally distinct fibril polymorphs with differential cytotoxicity, seeding properties, and resistance to PK digestion [57]. More recently, it has been shown that α-syn PFFs can contain of mixtures of structural assemblies that differ in the relative positioning of the C-terminal fibril fuzzy coat domain [46]. Similar to what we observed in M83^+/+^ mice, transgenic mice overexpressing P301L-mutant human tau develop variable signatures of misfolded tau, which has been attributed to stochastic generation of different tau strains [58, 59]. Moreover, recombinant prion protein can spontaneously misfold into several different self-propagating conformations under identical conditions [60, 61]. Therefore, stochastic misfolding of proteins into distinct aggregate strains may constitute a general mechanism underlying disease heterogeneity across several human neurodegenerative disorders.

Although our experimental approach was designed to remove other variables that could influence α-syn strain generation, it is conceivable that stochastic strain formation may also be influenced by other factors, such as the cell type in which the aggregates first appear [48]. For example, due to the existence of unique cellular milieus between different cell types in the brain, the α-syn aggregate structure associated with MSA may be selectively stabilized if it initially forms stochastically in oligodendrocytes, whereas propagation of the α-syn aggregate type associated with PD may be favored if the aggregates are initially generated in neurons. However, since the α-syn pathology in the brains of spontaneously ill M83^+/+^ mice is predominantly neuronal regardless of disease subtype, this suggests that distinct α-syn strains can originate from a single cell type. It is also possible that different types of aggregates can form stochastically in a common cell type, such as a certain population of neurons, and then exhibit conformation-specific differences in their abilities to spread to other cell types. This could explain why, despite the predominance of oligodendroglial α-syn pathology, neuronal α-syn inclusions can also be found in MSA and, conversely, why lower amounts of astroglial and oligodendroglial α-syn pathology can be found in the brains of PD patients despite a neuron-dominant LB signature [62, 63]. α-Syn post-translational modifications (PTMs) could also conceivably modulate strain generation and propagation. α-Syn can undergo a diverse range of PTMs including serine and tyrosine phosphorylation, ubiquitination, nitration, glycosylation, and C-terminal truncation [64, 65]. While M83^+/+^_Y_ was the most frequently observed synucleinopathy subtype in spontaneously sick M83^+/+^ mice, neither A53T_1_ nor A53T_2_ PFFs induced this disease signature. One potential explanation for this paradox is that a specific PTM is required for formation of the M83^+/+^_Y_-associated α-syn strain *in vivo*. Because recombinant A53T-mutant α-syn lacks potentially relevant PTMs, such as phosphorylation, it may be unable to adopt the aggregate structure present in the brains of mice exhibiting the M83^+/+^_Y_ synucleinopathy subtype. Indeed, polymerization of recombinant α-syn phosphorylated at Ser129 induces formation of a distinct α-syn polymorph [66].

We found that A53T-mutant PFFs were less diverse (2 PFF polymorphs giving rise to 2 distinct outcomes in M83^+/-^ mice) compared to WT α-syn PFFs (6 PFF polymorphs giving rise to 4 distinct outcomes upon propagation in M83^+/-^ mice). This is consistent with the multitude of WT α-syn PFF polymorph structures that have been solved using cryo-EM [67–71], whereas only three A53T-mutant α-syn PFF structures have been deposited in the Protein Data Bank to date (6LRQ, 7WNZ, and 7WO0) [72]. Therefore, mutations in α-syn may reduce the conformational spectrum of aggregates that can be formed. This provides a potential explanation for why WT α-syn can misfold in several different ways to yield either PD or MSA whereas specific α-syn mutations, as are present in several genetic synucleinopathies, are associated with variable Parkinsonian phenotypes but not with MSA [73, 74]. However, consistent with what we observed with A53T-mutant PFFs and M83^+/+^ mice, the A53T mutation can give rise to phenotypic variability among affected families [75–78].

Although we do not know whether the structures of the α-syn PFFs are conserved upon propagation in M83^+/-^ mice, different mechanisms may contribute to the growth and spread of α-syn aggregates *in vivo*. Fibrils can elongate by addition of α-syn monomers to either end via templated conformational conversion, resulting in structural fidelity and maintenance of strain properties [79]. In contrast, during secondary nucleation the surface of a fibril acts as a nucleation site that results in the growth of a new aggregate with a structure and strain properties that do not necessarily match those of the original fibril [80, 81]. Because the properties of the M83^+/+^ subtypes were maintained upon serial propagation, we speculate that these brain-derived α-syn strains propagate via templated conformational conversion. The consistent phenotypic differences in mice injected with replicates of A53T_1_ vs A53T_2_ PFFs is also suggestive of propagation via conformational templating. In contrast, at least a subset of the WT PFF polymorphs may initiate α-syn aggregation upon inoculation into M83^+/-^ mice by a secondary nucleation mechanism. This is supported by the following observations: 1) WT_3_ and WT_4_ PFFs had similar molecular structures yet initiated very different outcomes in M83^+/-^ mice; and 2) WT_1_ and WT_4_ PFFs had distinct molecular structures yet induced an indistinguishable synucleinopathy in M83^+/-^ mice. It is conceivable that a sequence mismatch between the WT α-syn PFFs and A53T-mutant α-syn-expressed by M83^+/-^ mice reduces the efficiency of conformational templating and thus promotes secondary nucleation or other types of strain adaptation. However, it is also possible that certain PFF preparations may contain a mixture of structurally diverse α-syn aggregates that differentially propagate and adapt upon inoculation into M83^+/-^ mice [82]. This could explain the observed discrepancies between WT PFF polymorph structure and the resultant synucleinopathy they cause as well as the discordant structural and propagation properties of the two replicates of WT_5_ PFFs.

Although structural diversity within α-syn aggregates clearly exists between individual PFF preparations and M83^+/+^ mice, there are several limitations to our study. As a proof-of-principle, we only investigated one α-syn concentration and one buffer condition when generating WT and A53T-mutant PFFs. It is known that different α-syn polymorphs or strains can arise when polymerizing the protein using different buffer conditions [42, 43, 47, 70] or in the presence of cofactors such as lipids or heparin [83, 84], and it is conceivable that fibrilization of recombinant α-syn at a higher concentration [85] could modulate the diversity of polymorphs that can form. Another limitation is that we only used M83 transgenic mice for the spontaneous disease and α-syn propagation experiments. As these mice overexpress mutant human α-syn using a non-native promoter, we do not know whether the α-syn strains present in the brains of inoculated M83^+/-^ mice have direct relevance to human disease-specific α-syn strains and whether the true strain diversity present in WT PFFs was underestimated due to an α-syn sequence mismatch between the PFFs and the mice. It should be noted that none of the WT or A53T-mutant PFF structures we solved using cryo-EM entirely match those obtained using α-syn aggregates purified from the brains of PD or MSA patients [38, 39]. In general, the structures of α-syn fibrils generated *in vitro* by spontaneous or seeded aggregation are not identical to those present in human brains [52, 86–91], and it has been argued that α-syn PFFs are more useful for studying general synucleinopathy mechanisms rather than being a precise model of PD [92]. Although we were unable to solve the structure of the rare WT_6_ PFF polymorph, it induced a similar synucleinopathy in M83^+/-^ mice to that observed following inoculation with brain extract from PD patients [53, 54], suggesting that potentially PD-relevant α-syn aggregates may emerge stochastically, albeit infrequently, during PFF formation.

The fact that considerable sample-to-sample structural and functional heterogeneity exists within α-syn aggregates has important practical implications. Our findings suggest that the inherent variability of PFFs and M83^+/+^ mice must be considered when designing and interpreting experiments that utilize these research tools. Thorough conformational characterization of individual α-syn PFF preparations should be undertaken prior to commencing experiments to ensure that the same polymorph is used for answering the research question of interest. Furthermore, *in vitro* methods for propagating individual PFF polymorphs with high conformational fidelity need to be developed so that large batches of conformationally homogeneous PFFs can be obtained. While using brain-derived α-syn seeds, such as those derived from MSA patients or MSA-inoculated M83^+/-^ mice [25, 26], may help to reduce experimental variability, this may not be possible in all situations and, despite their drawbacks, the simplicity and flexibility of α-syn PFFs make them ideal for research purposes. Finally, understanding α-syn conformational strains is likely to be crucial for the proper interpretation of therapeutic studies that utilize M83^+/+^ mice. Post hoc analysis of M83^+/+^ disease subtypes needs to be conducted to determine whether drugs targeting α-syn aggregation or propagation exhibit strain-specific effects. Alternatively, M83^+/-^ mice inoculated with the three M83^+/+^ subtypes could be used, which would facilitate the evaluation of α-syn strain-specific drug activity by removing the confounding factor of stochastic strain formation.

## Materials and Methods

### Expression and purification of recombinant α-syn

The coding sequence for untagged full-length WT or A53T-mutant human α-syn (residues 1-140) was inserted into the pET-28a vector (EMD Millipore), then expressed and purified from *E. coli* strain BL21(DE3). Briefly, the bacteria were treated with isopropyl β-d-1-thiogalactopyranoside (IPTG) for 2.5 h to induce α-syn expression, then collected by centrifugation at 5,000x *g* for 15 min. Cell pellets were washed in Dulbecco’s PBS (DPBS) (Gibco, #14190144), centrifuged again, and then frozen at -80 °C. Frozen cell pellets were resuspended in DPBS containing 1 mM phenylmethylsulfonyl fluoride (PMSF), lysed with a tip sonicator (6 pulses of 25 s each at 18% amplitude, with 2 min rest on ice after each pulse), and incubated in a boiling water bath for 15 min. Cellular debris was removed by centrifugation at 10,000x *g* at 4 °C for 10 min. The supernatant was then clarified by ultracentrifugation at 150,000x *g* at 4 °C for 30 min and dialyzed against Purification Buffer A [50 mM Tris-HCl, pH 8.3]. Recombinant α-syn was then subjected to fast protein liquid chromatography using a Mono Q anion exchange column (GE Healthcare), and α-syn was eluted with a linear gradient of 0 to 500 mM NaCl in Buffer A. Fractions were analyzed by SDS-PAGE and Coomassie blue staining, and those containing sufficiently pure α-syn were pooled and dialyzed against Purification Buffer B [20 mM Tris-HCl, pH 7.4]. After measuring sample absorbance at 280 nm using a NanoDrop spectrophotometer, the concentration of recombinant α-syn was calculated using Beer’s law and an extinction coefficient of 5,960 M^-1^cm^-1^. Purified protein was then flash-frozen in single-use aliquots and stored at -80 °C.

### Generation of α-syn PFFs

Purified recombinant α-syn, either WT or A53T-mutant, was polymerized in the presence of salt as previously described [47]. Briefly, 100 [47] mM NaCl was added to 1 mg/mL (69 µM) recombinant α-syn diluted in Purification Buffer B for a total volume of 400 µL, and the mixture was incubated with continuous shaking at 600 r.p.m. at 37 °C on an Eppendorf Thermomixer F1.5 for 7 days. Fibril formation was assessed by mixing polymerized α-syn (final concentration: 50 µg/mL) with 10 µM ThT (Sigma-Aldrich, #T3516) then measuring fluorescence using excitation and emission wavelengths of 444 nm and 485 nm, respectively. PFF preparations that exhibited a ThT fluorescence value of less than 1 x 10^6^ RFU (∼5-10 times that of a buffer-only blank) after 7 days incubation were put back onto the shaker for an additional 7 days (14 days total) with confirmation of ThT fluorescence at day 14.

### Negative stain electron microscopy

For each α-syn PFF sample, 2 µL of fibril solution were applied onto a freshly glow-discharged carbon-coated copper grid (EM Sciences, #ECF300-Cu). After 2 min incubation, the sample was carefully blotted off with filter paper. Next, 2 µL of 1% (w/v) uranyl acetate was applied on the top of the grid. After 1 min incubation, excess uranyl acetate was removed with filter paper and the grid was air-dried. Transmission electron microscopy images were acquired using a ThermoFisher Scientific Talos 120C instrument at an acceleration voltage of 120 kV. Images were collected using a 4k x 4k Ceta 16M CEMOS camera.

### Proteinase K digestion of PFFs

α-Syn PFFs were diluted to 50 µg/mL in 1X detergent buffer [0.5% (v/v) Nonidet P-40, 0.5% (w/v) sodium deoxycholate in DPBS] containing 50 µg/mL PK (Thermo Scientific, #EO0491) for a 1:1 α-syn:PK mass ratio. The sample was incubated with continuous shaking at 600 r.p.m. at 37 °C on an Eppendorf Thermomixer F1.5 for 1 h. After adding PMSF to a final concentration of 4 mM to inactivate PK, the insoluble fraction was collected by ultracentrifugation in a TLA-55 rotor (Beckman Coulter) at 100,000x *g* at 4 °C for 1 h. The pellet was resuspended in loading buffer [1X Bolt lithium dodecyl sulfate sample buffer (ThermoFisher Scientific, #B0007) containing 2.5% β-mercaptoethanol] and heated to 95 °C for 10 min. PK-digested samples were analyzed by SDS-PAGE, as described below, with detection of protease-resistant α-syn fragments by silver staining using the Pierce Silver Stain Kit according to the manufacturer’s protocol (Thermo Scientific, #24612).

### SDS-PAGE and immunoblotting

Samples were prepared in loading buffer and run on 4-12%, 10%, or 12% Bolt Bis-Tris Plus gels (Thermo Scientific) at 165 V for 38 to 48 min, or until the dye front reached the bottom of the gel. For immunoblotting, proteins were transferred onto a 0.45 µm-pore polyvinylidene fluoride membrane in transfer buffer [25 mM Tris-HCl, pH 8.3, 0.192 M glycine, 20% (v/v) methanol] at 35 V for 1 h. Proteins were crosslinked to the membrane in 0.4% (v/v) paraformaldehyde (Electron Microscopy Services, #15711) in DPBS by gentle rocking at 22 °C for 30 min, then rinsed twice with TBST [1X TBS containing 0.05% (v/v) Tween-20]. The membrane was then incubated in blocking buffer [5% (w/v) skim milk in TBST] at 22 °C for 1 h, then rocked in primary antibody (diluted in blocking buffer) at 4 °C overnight. The following primary antibodies were used in this study: anti-total α-syn mouse monoclonal Syn-1 (1:10,000 dilution; BD Biosciences, #610786), and anti-PSyn rabbit monoclonal EP1536Y (1:10,000 dilution; Abcam, #ab51253). Membranes were washed three times with TBST at 22 °C for 10 min each, then incubated at 22 °C for 1 h with horseradish peroxidase-conjugated secondary antibodies (Bio-Rad, #172-1011 or #172-1019) at 1:10,000 dilution in blocking buffer. After another three TBST washes, membranes were developed using Western Lightning enhanced chemiluminescence Pro (Revvity, #NEL122001EA) or SuperSignal West Dura Extended Duration Substrate (ThermoFisher Scientific, #37071), then either exposed to HyBlot CL X-ray film (Thomas Scientific, #1141J52) or imaged using the LiCor Odyssey Fc system. For quantitative comparison of protein signal across samples, equal amounts of each sample were loaded in each lane. For qualitative comparison of protease-resistant α-syn species, the amount of sample loaded was empirically adjusted by diluting in loading buffer so that banding pattern differences between samples could be readily visualized.

### Curcumin dye-binding assay

The fluorescent emission spectra for α-Syn PFFs following reaction with curcumin were obtained as previously described [47]. Briefly, 10 µM total α-syn (either PFFs or monomeric α-synuclein) was mixed with 5 µM curcumin (Sigma-Aldrich, #C1386) in a total volume of 100 µL. The sample was shaken at 850 r.p.m. at 22 °C for 15 min, and then dialyzed against dH_2_O for 50 min using a Slide-A-Lyzer™ MINI Dialysis Device with a 20-kDa molecular weight cut-off (ThermoFisher Scientific, #69590) to remove unbound dye. Dialyzed samples were pipetted in triplicate into black 96-well clear-bottom half-area plates (Greiner, #675096) and fluorescent emission spectra (460 ± 5 nm to 625 ± 5 nm) measured using a BMG CLARIOstar microplate reader after excitation at 432 ± 7.5 nm, with the gain set at 2,000. The background fluorescence was subtracted using a curcumin-only sample that was processed identically. Blank-corrected fluorescence values across replicates were averaged and then normalized to the highest signal within the spectrum, which was set at 1.0.

### Mice

Homozygous M83 (M83^+/+^) mice (stock number: 004479), which use the mouse prion protein promoter to overexpress human A53T mutant α-synuclein on a mixed C57BL/6 and C3H background [18], and non-transgenic B6C3F1 mice (stock number: 100010) were purchased from The Jackson Laboratory. They were then intercrossed to create hemizygous M83 (M83^+/-^) mice for inoculation experiments. The M83^+/+^ mice used for spontaneous disease studies were either purchased from The Jackson Laboratory or bred in-house. Mice were housed in single-sex groups of up to five animals per cage on a 12 h light/12 h dark cycle and had unlimited access to food and water. Both male and female mice were used in all experiments. All mouse experiments were done in accordance with guidelines set by the Canadian Council on Animal Care, under a protocol (AUP 4363.21) approved by the University Health Network Animal Care Committee.

### Intracerebral inoculations

For α-syn PFF injections, 100 µL aliquots were pipetted into thin-walled PCR microtubes and placed in a microtube holder (QSonica, #444), with the bottoms of the tubes suspended ∼1 cm high in a microplate horn sonicator (QSonica, #431MPX) attached to a circulating 37 °C water bath (QSonica, #4900-110). Samples were sonicated four times in 15 s pulses at 70% amplitude, with 2 min rest in between. Prior to inoculation, PFFs (or α-syn monomers) were diluted three-fold into inoculum diluent buffer [sterile PBS containing 5% (w/v) BSA]. For brain homogenate injections, stocks of frozen 10% (w/v) brain extract were thawed and diluted to 1% (w/v) using inoculum diluent buffer. Weanling M83^+/-^ mice at ∼5 weeks of age were injected non-stereotactically at a depth of ∼3 mm into the right cerebral hemisphere using a tuberculin syringe attached to a 27 gauge, 0.5-inch needle (BD Biosciences, #305945). Each mouse received 30 µL inoculum, which contained either 10 µg of recombinant α-syn (PFFs or monomeric α-syn) or 300 µg of homogenized brain, which corresponds to ∼30 µg total brain protein.

### Assessment of neurological disease and brain collection

Inoculated M83^+/-^ mice and uninoculated M83^+/+^ mice were monitored twice a week for signs of neurological illness such as hindlimb paralysis, bradykinesia, reduction in grip strength, and weight loss [47]. Observers were blind to the inoculum. When paralysis progressed to the point where mice exhibited a reduced ability to ambulate and obtain food, they were euthanized by perfusion with PBS and then their brains were removed and bisected parasagittally. The time between inoculation and euthanasia was recorded as the disease incubation period. The left hemisphere was flash-frozen on dry ice and stored at -80 °C for generation of brain homogenates. The right hemisphere was fixed in 10% (v/v) neutral buffered formalin for immunohistochemical analysis. If the mice did not develop neurological illness by 540 days post-inoculation, they were euthanized and their brains collected in the same manner. Mice found dead in their cages or euthanized due to non-synucleinopathy illnesses were excluded from the study (**Supplemental Tables 6-8**). Synucleinopathy was defined as the presence of detergent-insoluble and protease-resistant α-syn species in brain homogenates. To generate 10% (w/v) brain homogenates, frozen brain hemispheres were transferred into CK14 soft tissue homogenizing tubes containing 1.4 mm zirconium oxide beads (Bertin Technologies, #P000912-LYSK0-A). Following the addition of the appropriate volume of sterile PBS, tubes were placed in a Minilys homogenizer (Bertin Technologies) and homogenized using three 30 s pulses with 2-5 min rest in between each cycle. The resulting 10% (w/v) brain homogenates were then aliquoted and stored frozen at -80 °C.

### Detergent insolubility assays

Nine volumes of 10% brain homogenate were added to one volume of 10X detergent buffer [5% (v/v) Nonidet P-40, 5% (w/v) sodium deoxycholate in DPBS] containing Halt Phosphatase Inhibitor (ThermoFisher Scientific, #78420) and Pierce Universal Nuclease (ThermoFisher Scientific, #88701), and chilled on ice for 20 min. The mixture was subject to centrifugation at 1,500x *g* at 4 °C for 5 min to remove debris, and the protein concentration in the supernatant was determined by the bicinchoninic acid (BCA) assay (ThermoFisher Scientific, #23227). Samples were diluted to 1 mg/mL total protein concentration with 1X detergent buffer, chilled on ice for 20 min, and ultracentrifuged in a TLA-55 rotor at 100,000x *g* at 4 °C for 1 h. Pellets were resuspended in loading buffer and heated to 95 °C for 10 min, and insoluble PSyn levels were analyzed by SDS-PAGE and immunoblotting, then quantified by densitometry using ImageJ.

### Protease digestion assays on brain homogenates

Detergent-extracted brain homogenates were generated as described above and then samples were diluted to a concentration of 5 mg/mL total protein using 1X detergent buffer. Samples were digested with either 50 µg/mL thermolysin (TL) or 100 µg/mL proteinase K (PK), for a total protease/protein (w/w) ratio of 1:100 and 1:50, respectively. After shaking at 600 r.p.m. at 37 °C on an Eppendorf Thermomixer F1.5 for 1 h, digestions were stopped by adding EDTA to a final concentration of 5 mM (for TL digestions) or PMSF to a final concentration of 4 mM (for PK digestions). The insoluble fraction was collected by ultracentrifugation in a TLA-55 rotor at 100,000x *g* at 4 °C for 1 h, and pellets were resuspended in loading buffer and heated to 95 °C for 10 min. Protease-resistant α-syn or PSyn fragments were then visualized by SDS-PAGE and immunoblotting.

### Conformational stability assays

For analysis of α-syn PFFs, 5 µL of 1 mg/mL fibril preparations were mixed with 15 µL of 1X detergent buffer and 20 µL of 2X guanidine hydrochloride (GdnHCl) stock to make a final GdnHCl concentration of either 0, 1.0, 1.5, 2.0, 2.5, 3.0, 3.5, or 4.0 M. For analysis of α-syn aggregates in brain homogenates, samples were first detergent-extracted and clarified as described above. Then, 20 µL of 2X guanidine hydrochloride (GdnHCl) stock were added to an equal volume of detergent-extracted 10% (w/v) brain homogenate to achieve final GdnHCl concentrations of 0, 1.0, 1.5, 2.0, 2.5, 3.0, 3.5, or 4.0 M. Samples were shaken at 800 r.p.m. at 22 °C for 2 h, and then the GdnHCl concentration in each tube was normalized to 0.4 M in a total volume of 500 µL 1X detergent buffer. Samples were then treated with 10 µg/mL TL to remove non-aggregates α-syn species. After shaking at 600 r.p.m. at 37 °C on an Eppendorf Thermomixer F1.5 for 1 h, TL digestion was stopped by adding EDTA to a final concentration of 1 mM. Mixtures were then ultracentrifuged in a TLA-55 rotor at 100,000x *g* at 4 °C for 1 h. The pellet was resuspended in loading buffer and heated to 95 °C for 10 min, and levels of residual insoluble α-syn species were analyzed by SDS-PAGE and immunoblotting. Densitometry was performed using ImageJ, and the values were normalized with the 0 M GdnHCl sample, which was set at 100%. The GdnHCl_50_ value was calculated using the sigmoidal dose-response (variable slope) equation in GraphPad Prism and fixing the top and bottom values at 100 and 0, respectively. Denaturation curves were generated by averaging the normalized signal values for each individual replicate.

### Immunohistochemistry

Formalin-fixed mouse hemibrains were processed and embedded in paraffin, then sectioned into 4 µm sagittal slices and adhered to Superfrost Plus glass slides (Fisher Scientific, #22037246). Slides were placed on a 60 °C heat block for 20 min, then deparaffinized and rehydrated in a graded series of xylenes and ethanol. Slides were submerged in citrate buffer [10 mM sodium citrate, pH 6, with 0.05% (v/v) Tween-20] and steamed at high pressure for 15 min in a pressure cooker (Instant Pot, #112-0141-01) for epitope retrieval. They were then cooled to 22 °C and washed under running tap water for 10 min. To quench endogenous peroxidase activity, the slides were gently shaken in 3% H_2_O_2_ in PBS for 5 min then washed three times, 5 min each, with PBST [PBS containing 0.1% (v/v) Tween-20]. Slides were blocked with 2.5% (v/v) normal horse serum (Vector Laboratories, #MP-7401) at 22 °C for 1 h, then stained for PSyn using the antibody EP1536Y (Abcam, #ab51253) diluted 1:320,000 in DAKO antibody diluent (Agilent, #S0809) at 4 °C overnight. Slides were washed three times with PBST, 5 min each, then incubated in secondary antibody [ImmPress horseradish-peroxidase-labeled horse anti-rabbit detection kit (Vector Laboratories, #MP-7401)] at 22 °C for 30 min. After three 5-min washes in PBST, slides were developed for 1 min using ImmPACT 3,3’-diaminobenzidine peroxidase substrate (Vector Laboratories, #SK-4105) and rinsed under running tap water for 10 min. The slides were then counterstained with hematoxylin (Sigma-Aldrich, #GHS132) for 2 min, rinsed under running tap water for another 10 min, then dehydrated in a graded series of ethanol and xylenes. Slides were mounted using Cytoseal 60 (Epredia, #8310–4) then scanned with the TissueSnap and TissueScope LE120 systems (Huron Digital Pathology) and visualized with PMA.start (Pathomation).

The area covered by PSyn staining was quantified using ImageJ as previously described [47, 51], except the threshold range was adjusted to 0-100 for improved signal-to-noise ratio. Manual counting was used to determine the number of PSyn-positive neurons in the cerebral cortex, CA1, and dentate gyrus, the number of PSyn-positive astrocytes in the thalamus, as well as the morphology of PSyn-positive neurons in the midbrain.

### Cryo-EM grid preparation and imaging

For cryo-EM grid preparation, 3 µL of fibril solution was applied to freshly glow-discharged R1.2/1.3 holey carbon film grids (Quantifoil). After the grids were blotted for 6 s, the grids were flash-frozen in liquid ethane using a Mark IV Vitrobot (ThermoFisher Scientific) operated at 4 °C and 95% rH. Cryo-EM datasets were collected on a Talos Arctica (ThermoFisher Scientific) and a Titan Krios G4 transmission-electron microscope (ThermoFisher Scientific) and details are provided in **Supplemental Table 9**. Collected images were motion corrected and dose weighted on-the-fly using Warp [93].

### Helical reconstruction of α-syn fibrils

α-Syn fibrils were reconstructed using RELION-3.1 [94], following the helical reconstruction scheme [95]. First, the estimation of contrast transfer function parameters for each motion-corrected micrograph was performed using CTFFIND4 [96]. Next, filament picking was done using crYOLO [97]. For 2D classification, we extracted particle segments using a box size of 600 pixels downscaled to 200 pixels and an inter-box distance of 17 pixels. For 3D classification, the classified segments after 2D classification were re-extracted using a box size of 300 pixels or 320 pixels (**Supplemental Table 9**) without downscaling. Starting from featureless cylinder filtered to 60 Å, several rounds of refinements were performed while progressively increasing the reference model’s resolution. The helical rise was initially set to 4.75 Å and the twist was estimated from the micrographs. Once the β-strands were separated along the helical axis, we optimized the helical parameters (final parameters are reported in **Supplemental Table 9**). We then performed a gold-standard 3D auto-refinement, followed by standard RELION post-processing with a soft-edged solvent mask that includes the central 10% of the box height yielded post-processed maps (B-factors are reported in **Supplemental Table 9**). The resolution was estimated from the value of the Fourier shell correlation curve for two independently refined half-maps at 0.143 (**Supplemental Fig. 12**). The optimized helical geometry was then applied to the post-processed maps yielding the final maps used for model building.

### Atomic model building and refinement

For all α-syn fibrils, one protein chain was extracted from a previously resolved fibril structure (see **Supplemental Table 10** for details). Subsequent refinement in real space was conducted using PHENIX [98, 99] and Coot [100] in an iterative manner. The resulting models were validated with MolProbity [101].

### Statistical analysis

Statistical analyses were performed using GraphPad Prism (version 10.4.1) with a significance threshold of P < 0.05. Kaplan-Meier curves of mice following inoculation were compared using the Log-rank test. Normality of datasets was not assumed. For comparisons of two groups, a Mann-Whitney test was performed. For comparisons of three or more groups, a Kruskal-Wallis test followed by a Dunn’s multiple comparisons test was performed. To compare PSyn deposition in different brain regions, as well as the morphology of induced deposits, a two-way ANOVA followed by a Šídák’s multiple comparisons test was used.

## Acknowledgements

The authors thank Gabor Kovacs and Ali Karakani for assistance with the scanning of pathology slides. This work was funded by a grant to JCW from the Canadian Institutes of Health Research (PJT-169042). RWLS was supported by fellowships from the Croucher Foundation, Parkinson Canada, the Peterborough K.M. Hunter Charitable Foundation, as well as an Ontario Graduate Scholarship. BF was partially supported by a Mitacs Globalink Research Award. NRGS was supported by a Parkinson Canada/Parkinson Society of British Columbia doctoral fellowship, a doctoral research award from the Canadian Institutes of Health Research, and an Ontario Graduate Scholarship. Experimental schematics were generated using BioRender.com.

## Availability of data and material

The atomic models have been deposited in the Protein Data Bank (PDB) and the corresponding cryo-EM maps have been deposited in the Electron Microscopy Data bank (EMDB). All accession numbers are reported in **Supplemental Table 10**. All other data generated or analyzed during this study are included in the manuscript or the supplemental information.

## Competing interests

BF is an employee of AstraZeneca. The other authors have no competing interests to declare that are relevant to the content of this manuscript.

## Author contributions

Conceptualization: RWLS, JCW; Funding Acquisition: JCW, GFS; Formal Analysis: RWLS, BF, JDC, GFS, JCW; Investigation: RWLS, BF, JDC, NRGS, AM, ES; Supervision: GFS, JCW; Visualization: RWLS, BF, JDC, JCW; Writing—Original Draft Preparation: RWLS, JCW; Writing—Review and Editing: RWLS, BF, JDC, NRGS, AM, ES, GFS, JCW.

