## Supplemental information for "Stochastic Misfolding Drives the Emergence of Distinct α-Synuclein Strains"

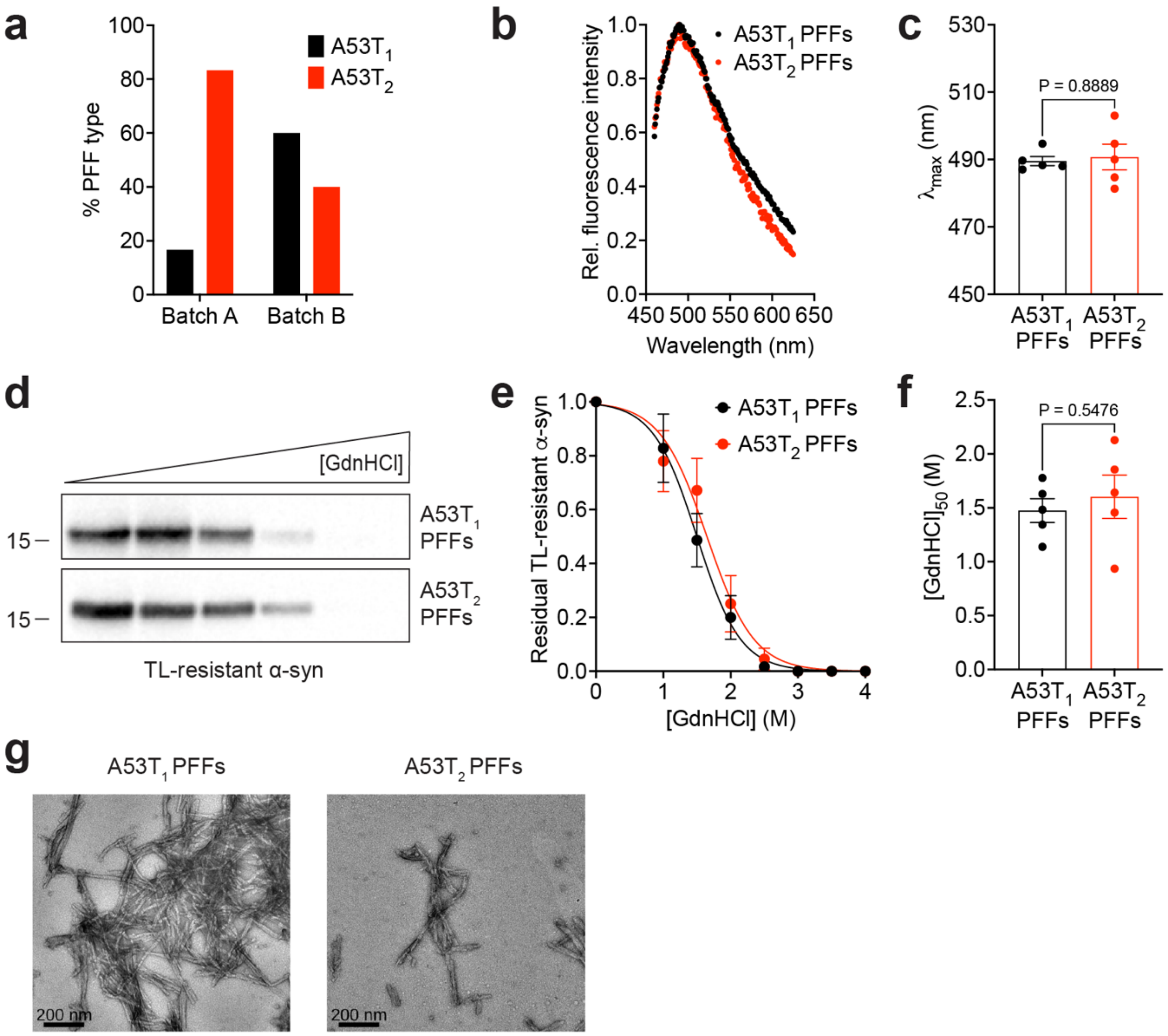


**Supplemental Figure 1. Additional characterization of A53T PFF polymorphs. a**) Percentage of A53T$_1$ and A53T$_2$ PFFs obtained from two independent batches of recombinant α-syn (Batch A: n = 12 PFFs; Batch B: n = 15 PFFs). **b**) Curcumin fluorescence emission spectra for A53T$_1$ and A53T$_2$ PFFs. The graph displays mean values for 5 independent PFF preparations for each polymorph. **c**) Peak curcumin emission spectra wavelengths ($\lambda_{max}$) for A53T$_1$ and A53T$_2$ PFFs (n = 5 each). Data is mean ± s.e.m. and statistical significance was assessed using a Mann-Whitney test. **d**) Representative immunoblots for residual insoluble TL-resistant α-syn species following treatment of A53T$_1$ and A53T$_2$ PFFs with increasing concentrations of GdnHCl. Molecular weight markers indicate kDa. **e**) Quantification of residual TL-resistant insoluble α-syn levels (mean ± s.e.m.) for A53T$_1$ and A53T$_2$ PFFs (n = 5 each) treated with increasing concentrations of GdnHCl. **f**) [GdnHCl]$_{50}$ values (mean ± s.e.m.) for A53T$_1$ and A53T$_2$ PFFs (n = 5 each). Statistical significance was assessed using a Mann-Whitney test. **g**) Representative negative stain electron microscopy images of A53T$_1$ and A53T$_2$ PFFs. Scale bars indicate 200 nm.

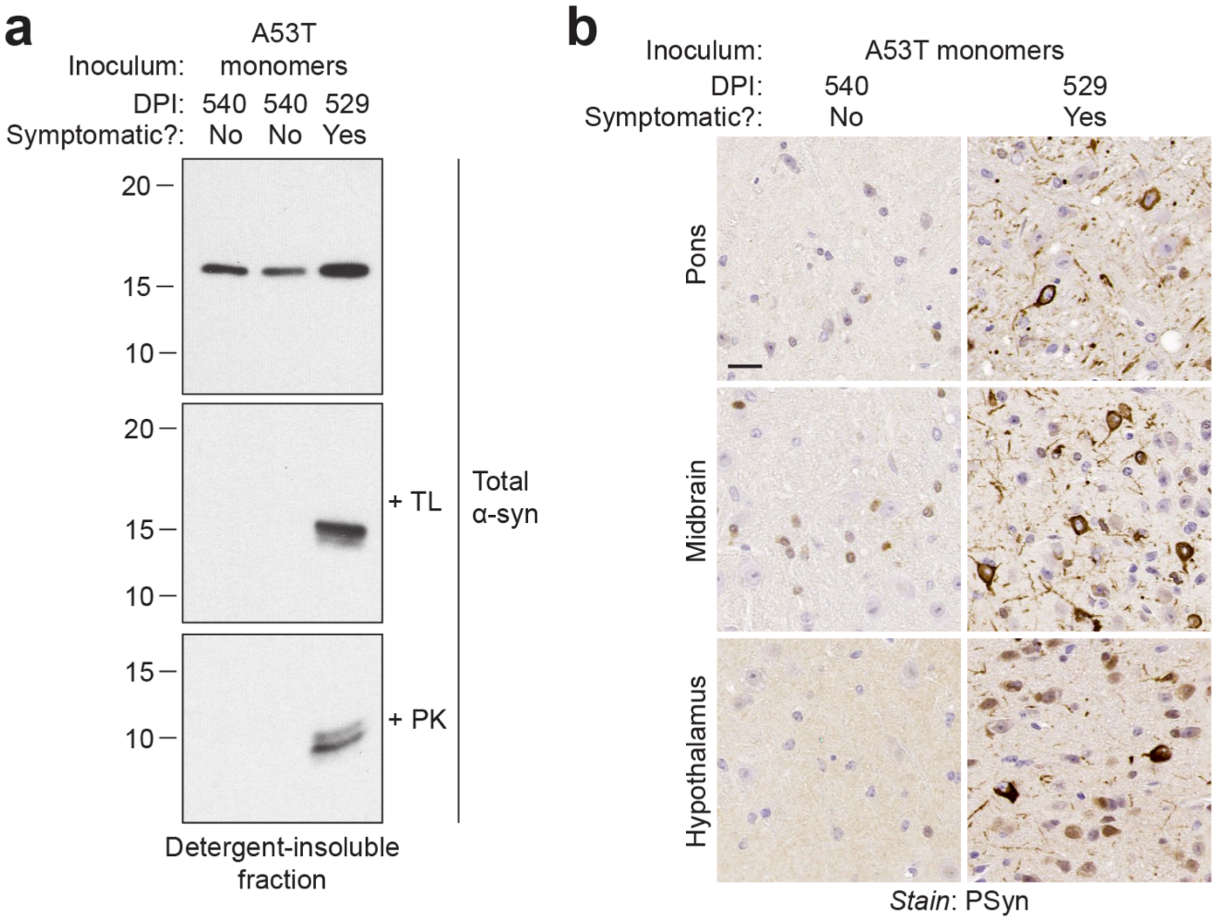


**Supplemental Figure 2. Characterization of M83$^{+/-}$ mice inoculated with monomeric A53T-mutant human α-syn. a**) Immunoblots of detergent-insoluble (top blot), TL-resistant insoluble (middle blot), and PK-resistant insoluble (bottom blot) α-syn species in brain extracts from M83$^{+/-}$ mice inoculated with monomeric A53T-mutant human α-syn. Brain samples from two asymptomatic mice at 540 days post-inoculation (DPI) and one symptomatic mouse at 529 DPI were analyzed. Molecular weight markers indicate kDa. **b**) Representative images of PSyn-stained sections of the pons, midbrain, and hypothalamus from M83$^{+/-}$ mice inoculated with monomeric A53T-mutant human α-syn. Brains from an asymptomatic mouse at 540 DPI and a symptomatic mouse at 529 DPI are shown. Scale bar indicates 20 μm (applies to all images). The protease digestion and neuropathological experiments suggest that the single symptomatic M83$^{+/-}$ mouse obtained following inoculation with monomeric A53T-mutant human α-syn exhibits a synucleinopathy signature consistent with the M83$^{+/+}{}_{X}$ subtype.

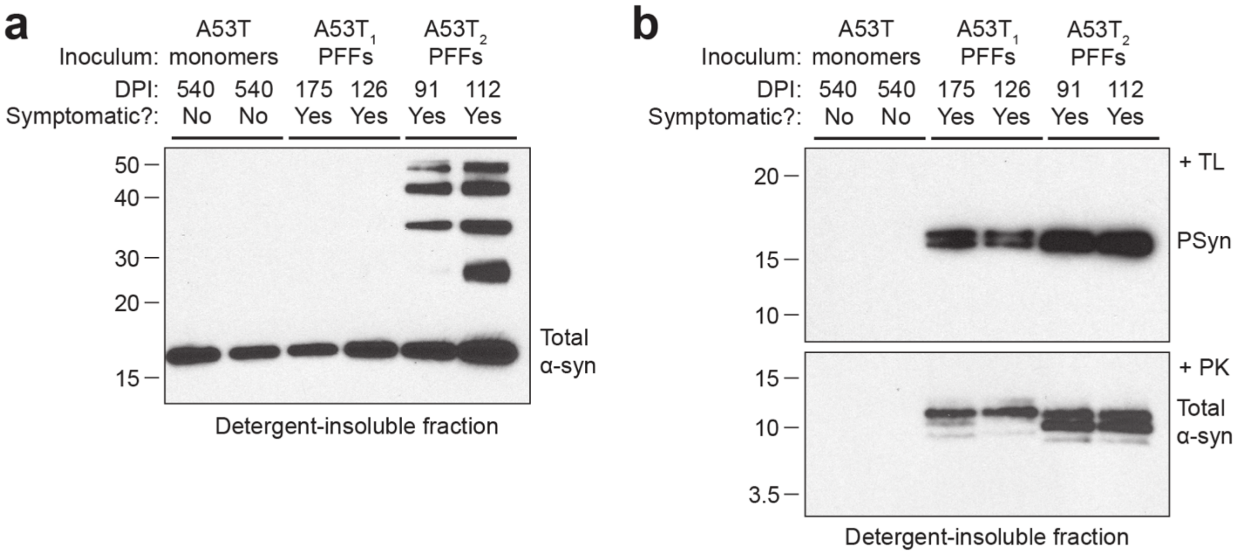


**Supplemental Figure 3. Additional biochemical characterization of M83$^{+/-}$ mice inoculated with A53T$_1$ or A53T$_2$ PFFs. a**) Immunoblot of detergent-insoluble total α-syn in brain extracts from M83$^{+/-}$ mice at the indicated DPI with either monomeric A53T α-syn (asymptomatic mice) or A53T PFFs (symptomatic mice). Two representative mice for each inoculum are shown. **b**) Immunoblots of detergent-insoluble TL-resistant PSyn species and PK-resistant total α-syn species in brain extracts from M83$^{+/-}$ mice at the indicated DPI with either monomeric A53T α-syn (asymptomatic mice) or A53T PFFs (symptomatic mice). Two representative mice for each inoculum are shown. Molecular weight markers indicate kDa.

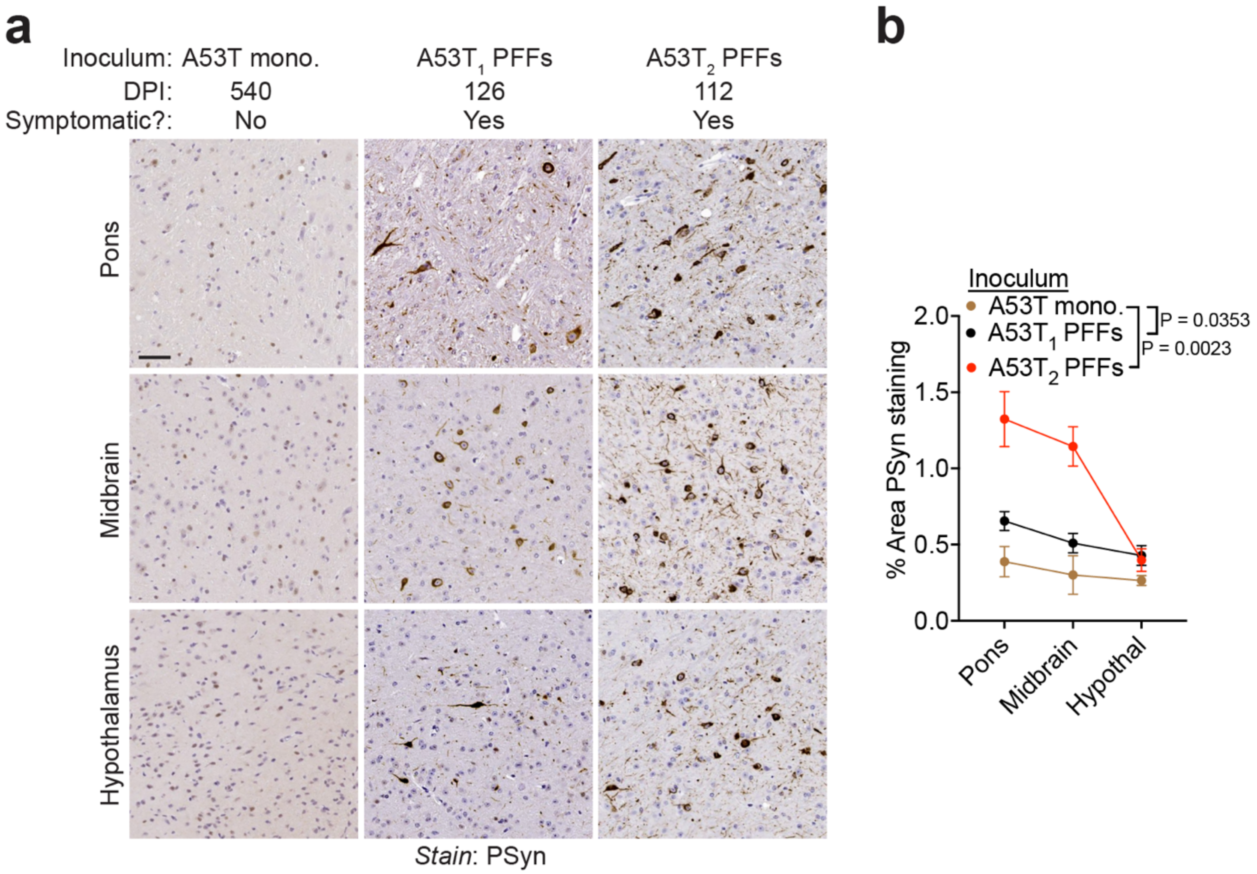


**Supplemental Figure 4. Induction of cerebral α-syn pathology in M83$^{+/-}$ mice inoculated with A53T PFFs. a**) Representative images of PSyn-stained sections of the pons, midbrain, and hypothalamus from an asymptomatic M83$^{+/-}$ mouse at 540 DPI with monomeric A53T-mutant human α-syn and symptomatic M83$^{+/-}$ mice at the indicated DPI with either A53T$_1$ or A53T$_2$ PFFs. Scale bar indicates 50 µm (applies to all images). **b**) Quantification of the area covered by PSyn staining (mean ± s.e.m) in the indicated brain regions from M83$^{+/-}$ mice injected with A53T monomers (n = 9), A53T$_1$ PFFs (n = 23), or A53T$_2$ PFFs (n = 25). All inoculated mice were included in the analysis, including A53T$_1$ PFF-inoculated mice that remained asymptomatic and the one monomer-injected mouse that became symptomatic. Statistical significance was assessed by two-way ANOVA.

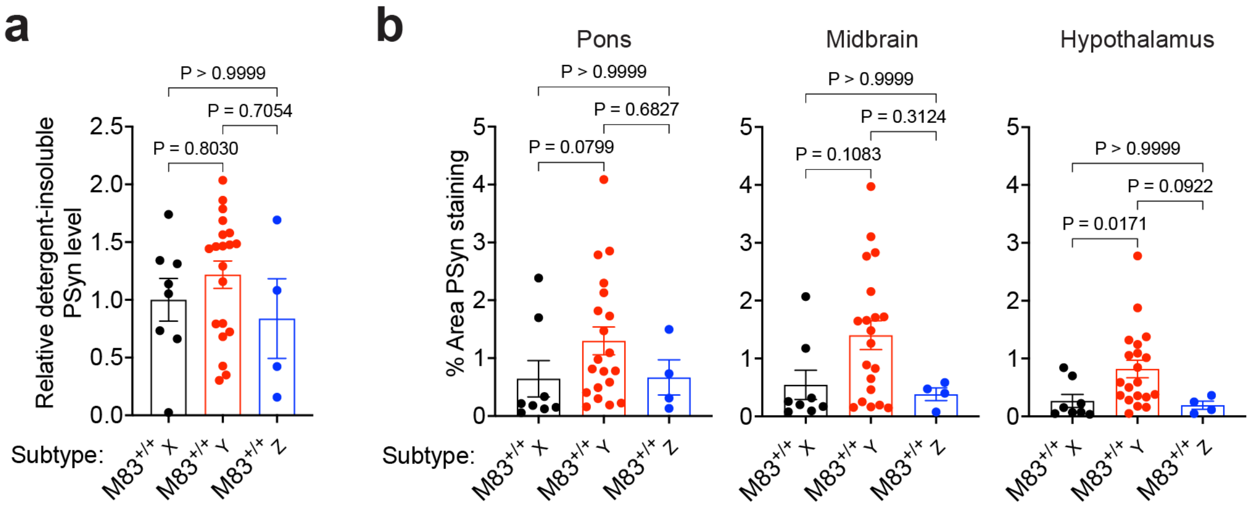


**Supplemental Figure 5. Relative amounts of PSyn deposition in spontaneously ill M83$^{+/+}$ mice exhibiting the X, Y, or Z molecular subtypes. a**) Quantification of detergent-insoluble PSyn species in brain extracts from symptomatic M83$^{+/+}$ mice classified as either subtype X (n = 8), subtype Y (n = 20), or subtype Z (n = 4). **b**) Quantification of the area covered by PSyn staining in the indicated brain regions from spontaneously ill M83$^{+/+}$ mice classified as either subtype X (n = 8), subtype Y (n = 20), or subtype Z (n = 4). In both panels, the graphs display mean ± s.e.m. Statistical significance was assessed using a Kruskal-Wallis test with Dunn's multiple comparison test.

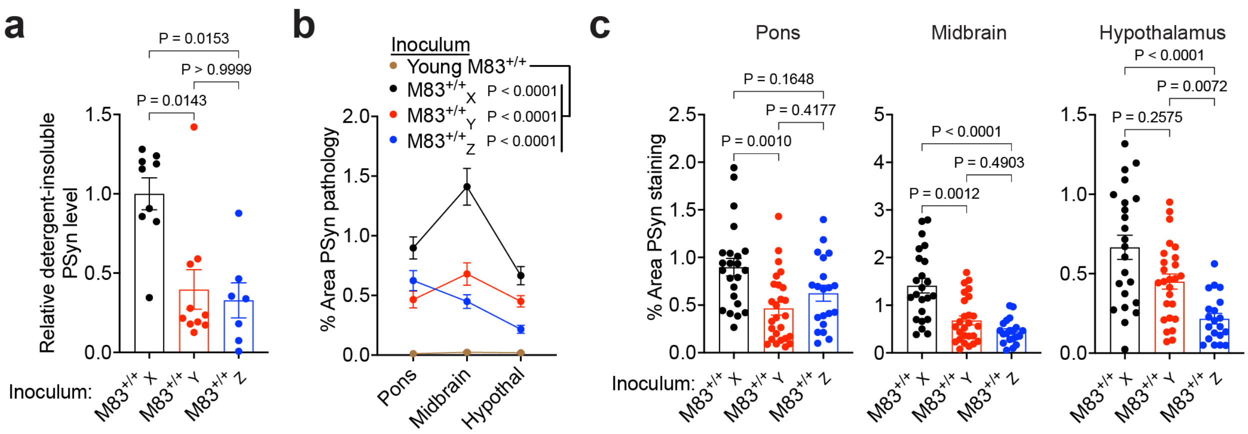


**Supplemental Figure 6. Relative amounts of PSyn deposition in M83$^{+/-}$ mice inoculated with M83$^{+/+}$ subtypes. a**) Quantification of detergent-insoluble PSyn species in brain extracts from symptomatic M83$^{+/-}$ mice inoculated with either M83$^{+/+}$ subtype X, Y, or Z (n = 9 each; 3 brains from each of 3 independent experiments per M83$^{+/+}$ subtype were analyzed). **b**) Comparison of the area covered by PSyn staining (mean ± s.e.m) in brains from asymptomatic M83$^{+/-}$ mice inoculated with young M83$^{+/+}$ brain extract (n = 8) to symptomatic M83$^{+/-}$ mice inoculated either the X (n = 23), Y (n = 26), or Z (n = 20) M83$^{+/+}$ subtype. **c**) Quantification of the area covered by PSyn staining in the indicated brain regions from symptomatic M83$^{+/-}$ mice inoculated with either M83$^{+/+}$ subtype X (n = 23), Y (n = 26), or Z (n = 20). In all panels, the graphs display mean ± s.e.m. In panels a and c, statistical significance was assessed using a Kruskal-Wallis test with Dunn's multiple comparison test. In panel b, statistical significance was assessed by two-way ANOVA.

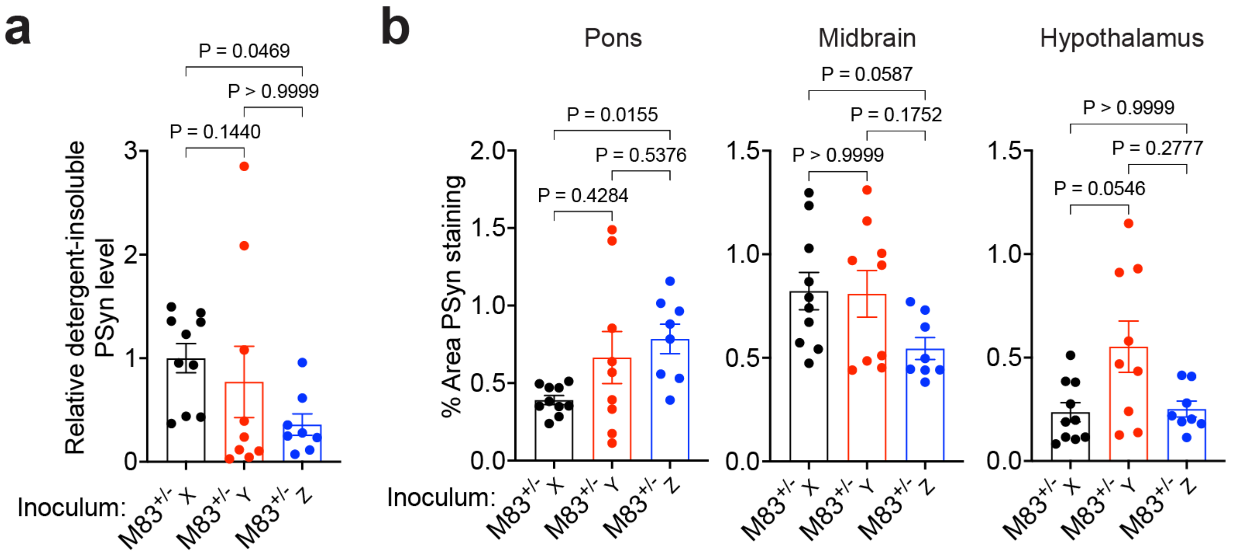


**Supplemental Figure 7. Relative amounts of PSyn deposition in M83$^{+/-}$ mice following serial passage of M83$^{+/+}$ subtypes. a**) Quantification of detergent-insoluble PSyn species in brain extracts from symptomatic M83$^{+/-}$ mice inoculated with either M83$^{+/-}$$_X$ (n = 10), M83$^{+/-}$$_Y$ (n = 9), or M83$^{+/-}$$_Z$ (n = 8) brain homogenate. **b**) Quantification of the area covered by PSyn staining in the indicated brain regions from symptomatic M83$^{+/-}$ mice inoculated with either M83$^{+/-}$$_X$ (n = 10), M83$^{+/-}$$_Y$ (n = 9), or M83$^{+/-}$$_Z$ (n = 8) brain homogenate. In both panels, the graphs display mean ± s.e.m. and statistical significance was assessed using a Kruskal-Wallis test with Dunn's multiple comparison test.

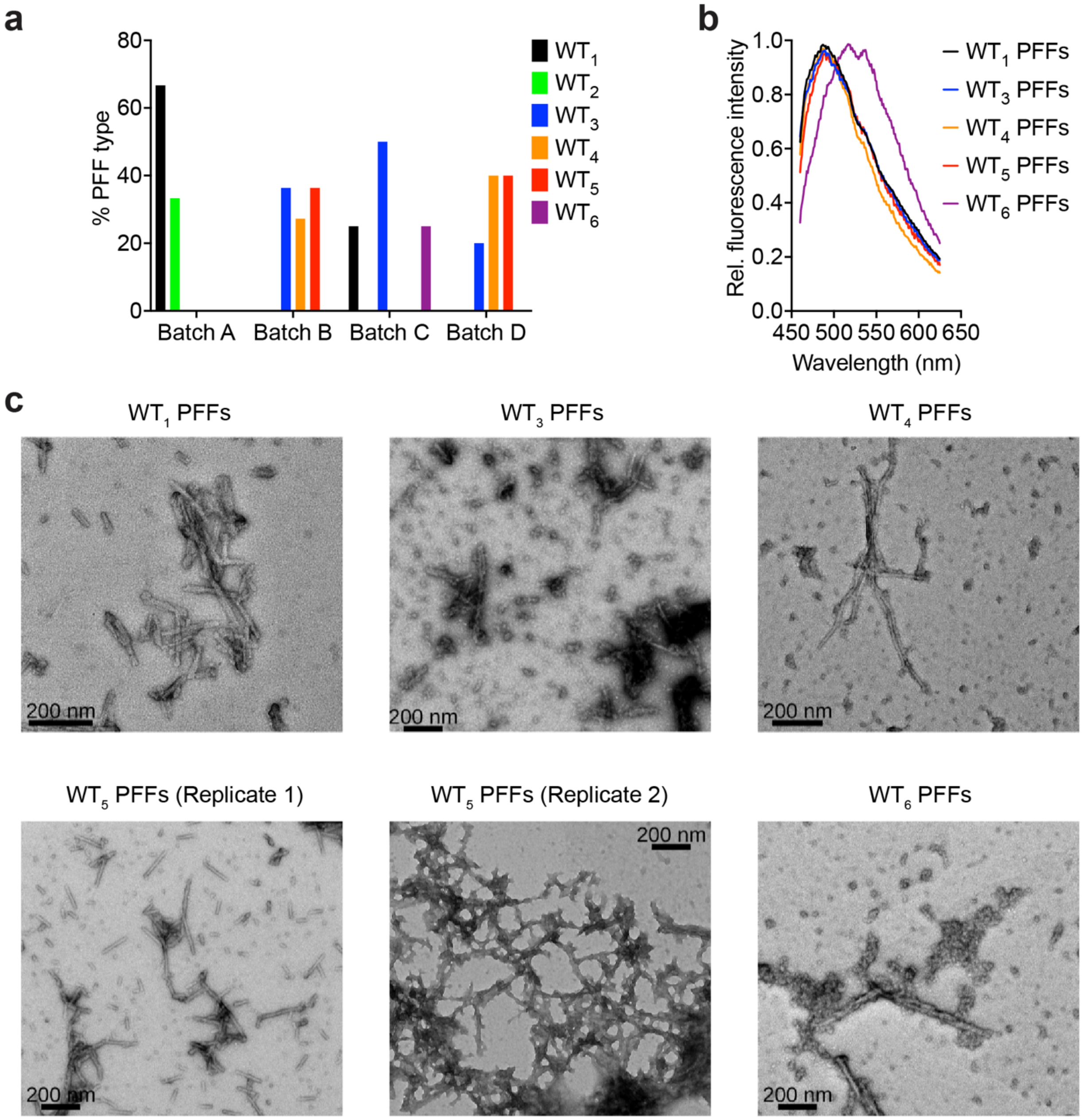


**Supplemental Figure 8. Additional characterization of WT PFF polymorphs. a**) Percentage of each WT PFF polymorph obtained from four independent batches of recombinant α-syn (Batch A: n = 3 PFFs; Batch B: n = 11 PFFs; Batch C: n = 4 PFFs; Batch D: n = 5 PFFs). **b**) Curcumin fluorescence emission spectra for $WT_1$ (n = 8), $WT_2$ (n = 1), $WT_3$ (n = 8), $WT_4$ (n = 7), $WT_5$ (n = 8), $WT_5$ (n = 10), and $WT_6$ (n = 1) PFF polymorphs. The graph displays mean values. **c)** Representative negative stain electron microscopy images of WT PFF polymorphs. Scale bars indicate 200 nm.

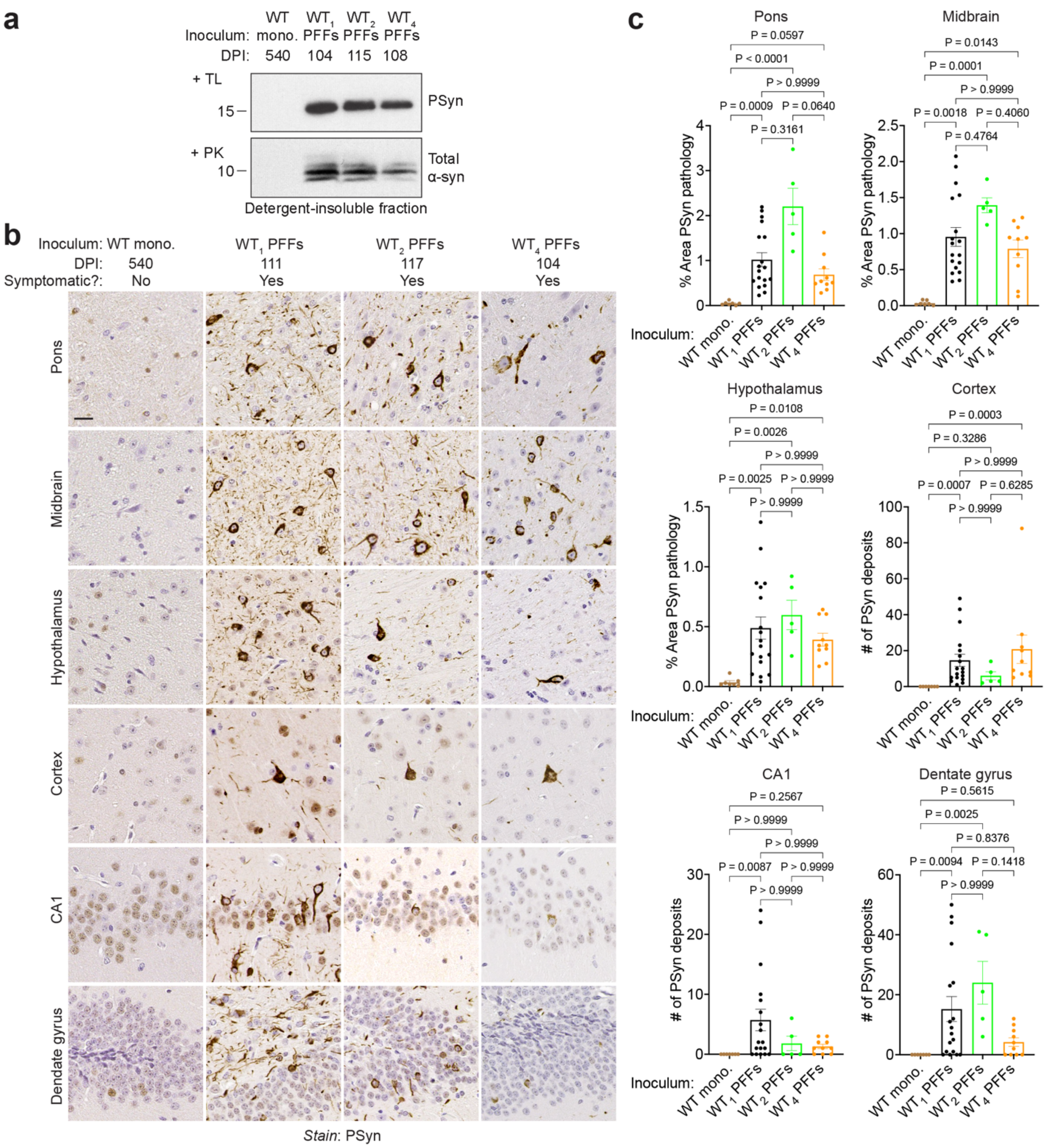


**Supplemental Figure 9. M83+/- mice inoculated with WT1, WT2, or WT4 PFFs develop a similar molecular phenotype. a**) Immunoblots of detergent-insoluble TL-resistant PSyn species and PK-resistant total α-syn species in brain extracts from symptomatic M83+/- mice at the indicated DPI with either WT1, WT2, or WT4 PFFs. Brain extract from an asymptomatic M83+/- mouse inoculated with monomeric (mono.) recombinant WT α-syn is included as a negative control. Molecular weight markers indicate kDa. **b**) Representative images of PSyn-stained sections of the pons, midbrain, hypothalamus, cortex, CA1, and dentate gyrus from an asymptomatic M83+/- mouse at 540 DPI with monomeric WT human α-syn and symptomatic M83+/- mice at the indicated DPI with either WT1, WT2, or WT4 PFFs. Scale bar indicates 20 µm (applies to all images). **c**) Quantification of the area covered by PSyn staining in the indicated brain regions from asymptomatic M83$^{+/-}$ mice at 540 DPI with monomeric α-syn (n = 7) and from symptomatic M83$^{+/-}$ mice inoculated with either $WT_1$ (n = 18), $WT_2$ (n = 5), or $WT_4$ (n = 10) PFFs. The graphs display mean ± s.e.m. and statistical significance was assessed using a Kruskal-Wallis test with Dunn's multiple comparison test.

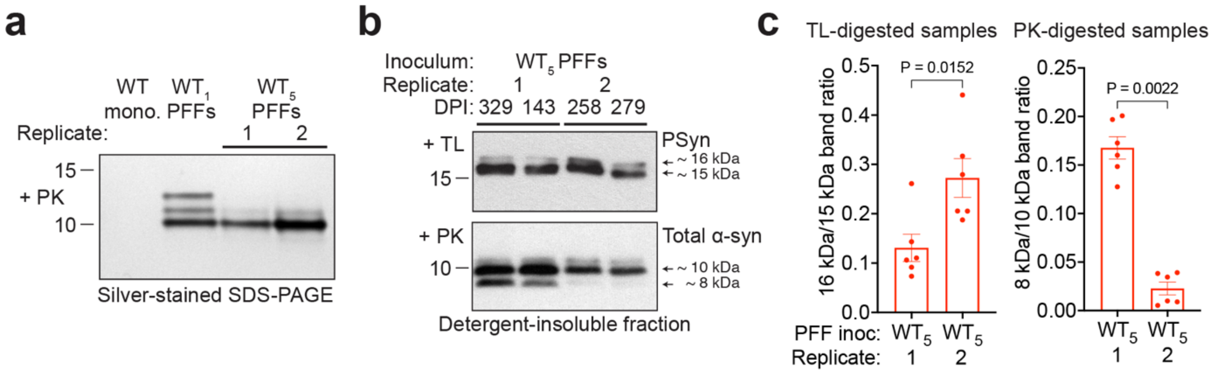


**Supplemental Figure 10. Divergent molecular phenotypes in M83$^{+/-}$ mice inoculated with different preparations of WT$_5$ PFFs. a**) Silver-stained SDS-PAGE of PK-resistant insoluble α-syn species in the preparations of WT$_1$ and WT$_5$ PFFs used for inoculation experiments in M83$^{+/-}$ mice. Unpolymerized monomeric (mono.) human WT α-syn is included as a negative control. **b**) Immunoblots of detergent-insoluble TL-resistant PSyn species and PK-resistant total α-syn species in brain extracts from symptomatic M83$^{+/-}$ mice at the indicated DPI with either of two different preparations of WT$_5$ PFFs. **c**) Quantification of the ratio of ~16 kDa to ~15 kDa TL-resistant species (left graph) and the ratio of ~8 kDa to ~10 kDa PK-resistant species (right graph) in brain extracts from symptomatic M83$^{+/-}$ mice inoculated with either of two distinct preparations of WT$_5$ PFFs (n = 6 each). The graphs display mean ± s.e.m. and statistical significance was assessed using a Mann-Whitney test. In panels a and b, molecular weight markers indicate kDa.

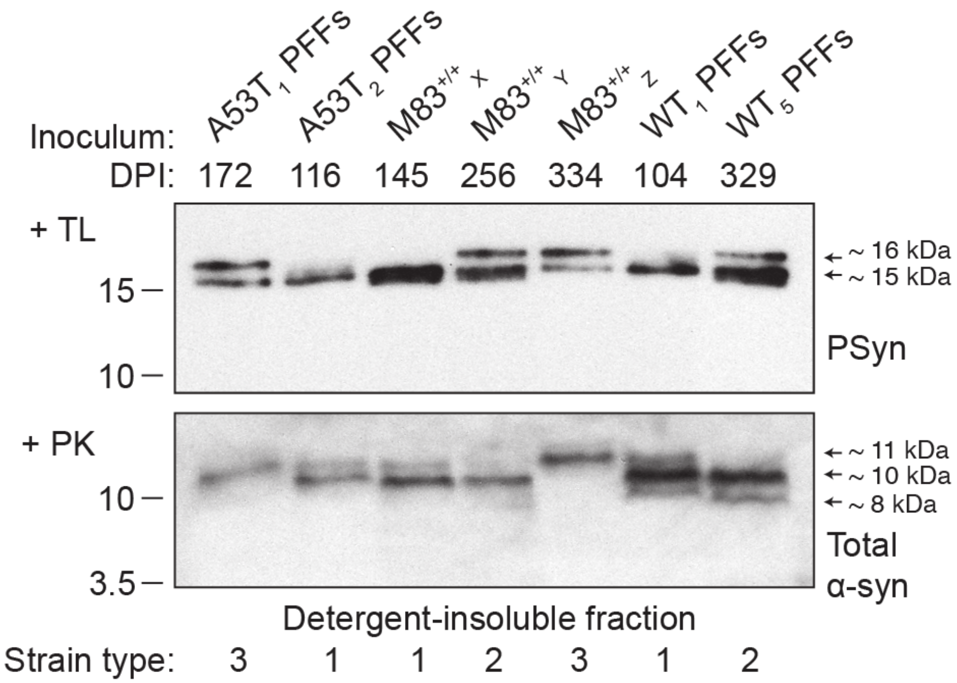


**Supplemental Figure 11. Molecular classification of α-syn strain types in $M83^{+/-}$ mice.** Immunoblots of detergent-insoluble TL-resistant PSyn species and PK-resistant total α-syn species in brain extracts from symptomatic $M83^{+/-}$ mice at the indicated DPI with the indicated preparations containing α-syn aggregates. Replicate 1 of $WT_5$ PFFs was used. Based on the banding pattern of TL- and PK-resistant α-syn species, three strain types were identified. Molecular weight markers indicate kDa.

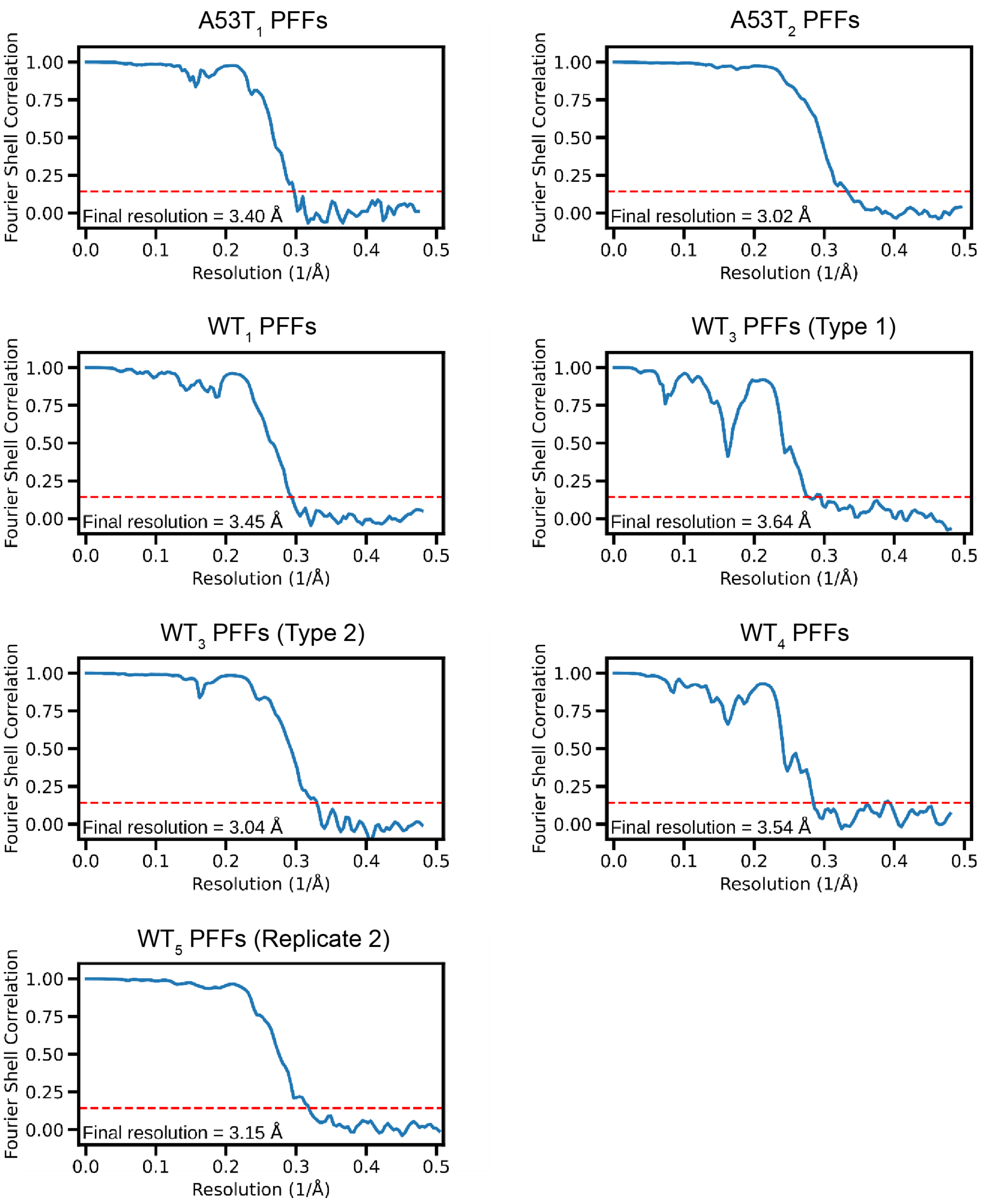


**Supplemental Figure 12. Fourier shell correlation (FSC) curves for A53T and WT PFF polymorphs.** Masked-corrected (z-percentage is 0.1) FSC curves. The final resolution is shown in the plot and was estimated from the value of the FSC curve for two separately refined masked half-maps at 0.143 (red line).

**Supplemental Table 1. Incubation periods for M83$^{+/-}$ mice inoculated with A53T PFFs**

| Inoculum | Replicate | PFF ThT fluorescence (RFU) | Incubation period (days ± s.e.m.) | n/n$_0$ |
|---|---|---|---|---|
| A53T monomers | - | - | > 540 | 1/9[1] |
| A53T$_1$ PFFs | 1 | 2.2 x 10$^7$ | 158 ± 7 | 8/8 |
| | 2 | 6.3 x 10$^6$ | 378 ± 42 | 7/8 |
| | 3 | 9.9 x 10$^6$ | 351 ± 6 | 3/7 |
| | Overall | - | 276 ± 30 | 18/23[2] |
| A53T$_2$ PFFs | 1 | 1.9 x 10$^6$ | 115 ± 4 | 8/8 |
| | 2 | 1.2 x 10$^6$ | 138 ± 2 | 9/9 |
| | 3 | 4.5 x 10$^6$ | 126 ± 1 | 8/8 |
| | Overall | - | 127 ± 2 | 25/25 |

n, number of mice with neurological illness; n$_0$, number of inoculated mice
[1]One mouse injected with A53T α-syn monomers developed neurological illness at 529 DPI
[2]Five mice remained healthy at 540 DPI

**Supplemental Table 2. Incubation periods for M83$^{+/-}$ mice inoculated with M83$^{+/+}$ brain extracts**

| M83$^{+/+}$ subtype | Replicate | Age of M83$^{+/+}$ mouse (days) | Incubation period (days ± s.e.m.) | n/n$_0$ |
|---|---|---|---|---|
| Young M83$^{+/+}$ | - | 62 | > 541 | 0/8 |
| M83$^{+/+}{}_X$ | 1 | 564 | 145 ± 5 | 9/9 |
| | 2 | 490 | 218 ± 26 | 8/8 |
| | 3 | 338 | 228 ± 22 | 6/6 |
| | Overall | - | 192 ± 13 | 23/23 |
| M83$^{+/+}{}_Y$ | 1 | 442 | 273 ± 8 | 10/10 |
| | 2 | 584 | 268 ± 15 | 8/8 |
| | 3 | 449 | 267 ± 6 | 8/8 |
| | Overall | - | 270 ± 6 | 26/26 |
| M83$^{+/+}{}_Z$ | 1 | 451 | 338 ± 25 | 7/7 |
| | 2 | 320 | 244 ± 17 | 6/6 |
| | 3 | 465 | 303 ± 18 | 7/9[1] |
| | Overall | - | 297 ± 14 | 20/22 |

n, number of mice with neurological illness; n$_0$, number of inoculated mice
[1]Two mice remained healthy at 540 DPI

**Supplemental Table 3. Incubation periods for second passage of M83$^{+/+}$ subtypes in M83$^{+/-}$ mice**

| Inoculum | Age of M83$^{+/-}$ mouse (days) | Incubation period (days ± SEM) | n/$n_0$ |
|---|---|---|---|
| M83$^{+/-}$$_X$ | 142 | 164 ± 14 | 10/10 |
| M83$^{+/-}$$_Y$ | 302 | 294 ± 23 | 9/9 |
| M83$^{+/-}$$_Z$ | 282 | 278 ± 24 | 8/8 |

n, number of mice with neurological illness; $n_0$, number of inoculated mice

**Supplemental Table 4. Incubation periods for M83$^{+/-}$ mice inoculated with WT α-syn PFFs**

| Inoculum | Replicate | PFF ThT fluorescence (RFU) | Incubation period (days ± SEM) | n/$n_0$ | Significance vs. monomer-injected mice[1] | Significance vs. $WT_5$ PFF-injected mice[1] |
|---|---|---|---|---|---|---|
| WT monomers | - | - | > 540 | 0/7 | - | P < 0.0001 |
| $WT_1$ PFFs | 1 | 1.6 x $10^7$ | 120 ± 3 | 9/9 | - | - |
| | 2 | 4.6 x $10^6$ | 132 ± 1 | 9/9 | - | - |
| | Overall | - | 126 ± 2 | 18/18 | P < 0.0001 | P < 0.0001 |
| $WT_2$ PFFs | - | 9.3 x $10^6$ | 122 ± 4 | 5/5 | P = 0.0003 | P < 0.0001 |
| $WT_3$ PFFs | - | 3.9 x $10^6$ | 455 | 1/8[2] | P = 0.3496 | P < 0.0001 |
| $WT_4$ PFFs | - | 1.2 x $10^6$ | 112 ± 3 | 10/10 | P < 0.0001 | P < 0.0001 |
| $WT_5$ PFFs | 1 | 2.5 x $10^6$ | 216 ± 33 | 6/6 | - | - |
| | 2 | 8.7 x $10^6$ | 242 ± 9 | 6/6 | - | - |
| | Overall | - | 229 ± 17 | 12/12 | P < 0.0001 | - |
| $WT_6$ PFFs | - | 1.2 x $10^6$ | 427 ± 44 | 2/6[3] | P = 0.1096 | P = 0.0001 |

n, number of mice with neurological illness; $n_0$, number of inoculated mice

[1]Statistical significance was assessed using the Log-rank test

[2]Seven mice remained healthy at 541 DPI

[3]Four mice remained healthy at 540 DPI

**Supplemental Table 5. Defining characteristics of α-syn strain types identified in the brains of inoculated M83$^{+/-}$ mice**

| Disease attribute | Strain type | | | |
|---|---|---|---|---|
| | 1 | 2 | 3 | 4 |
| Incubation period | 4-5 months | 7-10 months | 8-12 months | > 18 months |
| TL digestion pattern (PSyn) | Single dominant ~15 kDa band | Dominant ~15 kDa band with additional ~16 kDa band | Equal intensity ~15 and ~16 kDa bands | N/A |
| PK digestion pattern (total α-syn) | Dominant ~10 kDa band with additional ~11 kDa band | Dominant ~10 kDa band with additional ~8 kDa band | Single dominant ~11 kDa band | N/A |
| Morphology of PSyn deposits | Predominantly ring-like midbrain inclusions | Predominantly LB-like midbrain inclusions | Mixture of ring-like and LB-like midbrain inclusions | Sparse LB-like inclusions in periventricular region |
| Cortical PSyn deposition | Low | High | Low | None |
| *Example(s) from this study:* | $A53T_2$, $WT_1$, $WT_2$, and $WT_4$ PFFs; $M83^{+/+}{}_X$ | $WT_5$ PFFs (Replicate 1), $M83^{+/+}{}_Y$ | $A53T_1$ PFFs, $M83^{+/+}{}_Z$ | $WT_6$ PFFs |
| *Other potential examples:* | "S" strain; MSA-inoculated M83$^{+/-}$ mice | "NS" strain | | PD-inoculated M83$^{+/-}$ mice |

**Supplemental Table 6. List of M83$^{+/-}$ mice with intercurrent illness removed from the A53T PFF propagation studies**

| Inoculum | Sex | Days post-inoculation | Notes |
|---|---|---|---|
| A53T monomers | M | 127 | Found dead (likely due to fighting injury) |
| $A53T_1$ PFFs (Replicate 1) | M | 158 | Euthanized due to inability to urinate |
| $A53T_1$ PFFs (Replicate 3) | M | 423 | Found dead |
| $A53T_2$ PFFs (Replicate 1) | M | 43 | Found dead |
| | M | 105 | Euthanized due to penile injury |

**Supplemental Table 7. List of $M83^{+/-}$ mice with intercurrent illness removed from the $M83^{+/+}$ subtype serial propagation studies**

| Inoculum | Sex | Days post-inoculation | Notes |
|---|---|---|---|
| Young $M83^{+/+}$ | M | 73 | Found dead |
| | M | 467 | Found dead |
| $M83^{+/+}_{X}$ (Replicate 1) | M | 126 | Found dead |
| $M83^{+/+}_{X}$ (Replicate 2) | M | 231 | Euthanized due to atypical illness; no synucleinopathy found |
| | F | 418 | Euthanized due to atypical illness; no synucleinopathy found |
| $M83^{+/+}_{X}$ (Replicate 3) | M | 347 | Euthanized due to intestinal tumor |
| | F | 225 | Euthanized due to flooded cage |
| | F | 225 | Euthanized due to flooded cage |
| $M83^{+/+}_{Y}$ (Replicate 2) | M | 86 | Euthanized due to fighting injuries |
| | M | 243 | Found dead |
| $M83^{+/+}_{Y}$ (Replicate 3) | M | 230 | Found dead |
| $M83^{+/+}_{Z}$ (Replicate 1) | M | 289 | Found dead |
| | M | 352 | Euthanized due to fighting injuries |
| | F | 528 | Found dead |
| $M83^{+/+}_{Z}$ (Replicate 2) | M | 116 | Euthanized due to bladder stones |
| | M | 193 | Euthanized due to atypical illness; no synucleinopathy found |
| | M | 302 | Found dead |
| | F | 354 | Euthanized due to atypical illness; no synucleinopathy found |
| $M83^{+/+}_{Z}$ (Replicate 3) | M | 311 | Euthanized due to fighting injuries |
| $M83^{+/-}_{Y}$ | M | 222 | Euthanized due to fighting injuries |
| $M83^{+/-}_{Z}$ | M | 277 | Found dead |
| | F | 496 | Euthanized due to atypical illness; no synucleinopathy found |

**Supplemental Table 8. List of M83$^{+/-}$ mice with intercurrent illness removed from the WT PFF propagation studies**

| Inoculum | Sex | Days post-inoculation | Notes |
|---|---|---|---|
| WT monomers | M | 516 | Found dead |
| | M | 487 | Found dead |
| | F | 347 | Euthanized due to brain tumor |
| | F | 504 | Found dead |
| | F | 522 | Found dead |
| $WT_2$ PFFs | M | 119 | Found dead |
| | F | 63 | Found dead (flooded cage) |
| | F | 63 | Found dead (flooded cage) |
| | F | 63 | Found dead (flooded cage) |
| $WT_3$ PFFs | M | 86 | Found dead |
| | M | 237 | Euthanized due to atypical illness; no synucleinopathy found |
| $WT_5$ PFFs (Replicate 1) | M | 101 | Euthanized due to fighting injuries |
| | M | 101 | Euthanized due to fighting injuries |
| | M | 101 | Euthanized due to fighting injuries |
| | M | 111 | Found dead |
| $WT_5$ PFFs (Replicate 2) | M | 94 | Found dead |
| | M | 178 | Euthanized due to fighting injuries |
| $WT_6$ PFFs | M | 254 | Found dead |
| | M | 303 | Found dead |
| | M | 324 | Found dead |
| | M | 337 | Euthanized due to seizure |

### Supplemental Table 9. Cryo-EM data collection statistics for WT and A53T PFF polymorphs

| Data collection | PFF polymorphs | | | | | | |
|---|---|---|---|---|---|---|---|
| | A53T$_1$ | A53T$_2$ | WT$_1$ | WT$_3$ | | WT$_4$ | WT$_5$ (Replicate 2) |
| **Microscope** | Talos Artica | Titan Krios G4 | Titan Krios G4 | Titan Krios G4 | | Titan Krios G4 | Titan Krios G4 |
| **Voltage [keV]** | 200 | 300 | 300 | 300 | | 300 | 300 |
| **Detector** | K3 | K3 | Falcon 4 | Falcon 4 | | Falcon 4 | K3 |
| **Magnification** | 100,000 | 105,000 | 96,000 | 96,000 | | 96,000 | 105,000 |
| **Pixel size [Å]** | 0.8388 | 0.836 | 0.808 | 0.808 | | 0.808 | 0.82 |
| **Defocus range [µm]** | -0.5 to -2.5 | -0.5 to -2.5 | -0.5 to -2.5 | -0.5 to -2.5 | | -0.5 to -2.5 | -0.5 to -2.5 |
| **Exposure time [s/frame]** | 1.5 | 4.3 | 4.25 | 4.43 | | 4.22 | 1.4 |
| **Number of frames** | 30 | 35 | 1022 | 1043 | | 1015 | 40 |
| **Total dose [e$^-$/Å$^2$]** | ~30.6 | ~32.7 | ~40 | ~40 | | ~40 | ~40 |
| ***Reconstruction*** | | | | | | | |
| **Micrographs** | 14,343 | 11,445 | 8,543 | 9,120 | | 8,570 | 9,573 |
| **Box width [pixels]** | 300 | 300 | 320 | 320 | | 320 | 300 |
| **Inter-box distance [pixels]** | 17 | 17 | 17 | 17 | | 17 | 17 |
| **Picked segments (no.)** | 1,536,336 | 398,550 | 664,239 | 601,219 | | 3,193,361 | 1,557,543 |
| ***Final map*** | | | | Type 1 | Type 2 | | |
| **Final segments [no.]** | 37,465 | 15,954 | 24,963 | 5,076 | 20,107 | 8,212 | 38,375 |
| **Final resolution [Å] (FSC=0.143)** | 3.40 | 3.02 | 3.45 | 3.64 | 3,04 | 3.54 | 3.15 |
| **Applied map sharpening B-factor [Å$^2$]** | -100 | -97 | -64 | -40 | -53 | -53 | -116 |
| **Symmetry imposed** | C1 | C1 | C1 | C1 | C2 | C2 | C1 |
| **Helical rise [Å]** | 2.46 | 2.38 | 2.39 | 2.39 | 4.79 | 4.79 | 2.39 |
| **Helical twist [°]** | 179.76 | 179.31 | 179.65 | 179.37 | -1.5 | -1.5 | 179.45 |

### Supplemental Table 10. Cryo-EM model statistics for WT and A53T PFF polymorphs

| | PFF polymorphs | | | | | | |
|---|---|---|---|---|---|---|---|
| | A53T$_1$ | A53T$_2$ | WT$_1$ | WT$_3$ | | WT$_4$ | WT$_5$ (Replicate 2) |
| | | | | Type 1 | Type 2 | | |
| **PDB-ID** | 9rb3 | 9rb6 | 9rb7 | 9rb8 | 9rb9 | 9rba | 9rbb |
| **EMDB-ID** | EMD-53884 | EMD-53885 | EMD-53886 | EMD-53887 | EMD-53888 | EMD-53889 | EMD-53890 |
| **Model composition** | | | | | | | |
| Chains | 10 | 10 | 10 | 10 | 10 | 10 | 10 |
| Non-hydrogen atoms | 5200 | 4310 | 7340 | 3100 | 3100 | 6420 | 3160 |
| Protein residues | 750 | 620 | 530 | 440 | 440 | 440 | 470 |
| **RMS deviations** | | | | | | | |
| Bond lengths [Å] | < 0.01 | 0.02 | 0.02 | 0.02 | 0.02 | 0.02 | 0.02 |
| Bond angles [°] | 0.57 | 2.36 | 2.90 | 2.72 | 2.74 | 2.74 | 2.40 |
| **Validation** | | | | | | | |
| MolProbity score | 1.67 | 1.98 | 1.49 | 1.46 | 1.70 | 1.72 | 1.72 |
| Clash score | 4.42 | 1.13 | 9.26 | 5.95 | 6.10 | 6.42 | 5.43 |
| **Ramachandran plot** | | | | | | | |
| Outliers [%] | 0 | 0 | 0 | 0 | 0 | 0 | 0 |
| Allowed [%] | 7.04 | 6.67 | 1.96 | 0 | 0 | 0 | 2.22 |
| Favored [%] | 92.96 | 93.33 | 98.04 | 100 | 100 | 100 | 97.78 |
